# A multilevel formalism to model the hybrid E/M phenotypes in Epithelial-Mesenchymal Plasticity

**DOI:** 10.1101/2024.11.26.625479

**Authors:** Kishore Hari, Shubham Tripathi, Vaibhav Anand, Mohit Kumar Jolly, Herbert Levine

## Abstract

Epithelial–mesenchymal plasticity (EMP) is a cell-fate switching program that enables cells to adopt a spectrum of phenotypes ranging from epithelial (E) to mesenchymal (M), including intermediate hybrid E/M states. Hybrid E/M phenotypes are conducive to cancer metastasis, as they are associated with metastatic initiation, cancer stemness, drug resistance, and collective migration. Boolean models of the gene regulatory networks (GRNs) underlying EMP have yielded valuable insights into the dynamics of E and M phenotypes. However, these models are limited in their ability to capture hybrid phenotypes effectively, as they restrict gene expression to binary states. In contrast, hybrid E/M cells often exhibit partial expression of epithelial and mesenchymal markers. To overcome this limitation, we modified a threshold-based Boolean formalism to incorporate intermediate gene expression levels. The resulting multilevel model reveals novel hybrid steady-states characterized by partial expression of both E and M genes, thereby expanding the phenotypic landscape beyond that represented by traditional Boolean approaches. Notably, these hybrid states exhibit lower frustration compared to their counterparts in classical Boolean models. Furthermore, by resolving dynamical degeneracy, we demonstrate that the hybrid states identified by the multilevel model are more stable. These findings suggest that introducing minimal additional complexity into Boolean models can uncover previously hidden qualitative features of phenotypic landscapes governed by GRNs.

**Significance Statement:** Hybrid epithelial-mesenchymal phenotypes drive metastasis through collective migration, stemness, and drug resistance. However, traditional Boolean models of epithelial-mesenchymal plasticity (EMP) networks cannot capture the partial gene expression characteristic of these biologically crucial hybrid states. We developed a multilevel Boolean formalism that extends classical binary models to include intermediate expression levels. This approach successfully captures hybrid states with partial expression of both epithelial and mesenchymal genes, exhibiting lower frustration and greater stability than traditional Boolean hybrid phenotypes. While conventional Boolean hybrid states depend on modeling artifacts, multilevel hybrid states emerge independently of such uncertainties. Critically, plasticity experiments revealed that multilevel hybrid states show significantly higher transition probabilities to other hybrid states compared to traditional models, creating a stable “hybrid cloud” that maintains metastatic properties while allowing adaptive responses to environmental stress. This multilevel framework provides a biologically meaningful characterization of hybrid phenotypes essential for understanding metastasis.

## 1 Introduction

Metastasis, the leading cause of cancer mortality [1], is the process by which cancer cells disseminate from a primary tumor to multiple organs. Carcinomas, which originate from epithelial tissues, initially exhibit an epithelial phenotype characterized by strong cell-cell and cell-matrix adhesion. To metastasize, these cells must detach from the primary tumor, migrate through the bloodstream to a secondary site, and invade and colonize that site, often by adopting a mesenchymal phenotype. Subsequently, these cells may revert to an epithelial phenotype to establish secondary tumors [2]. This reversible transition between epithelial and mesenchymal states, termed epithelial-mesenchymal plasticity (EMP), is a critical driver of metastasis [3]. Among the various phenotypic states accessible through EMP, hybrid epithelial/mesenchymal (E/M) phenotypes, exhibiting both epithelial and mesenchymal traits, are highly conducive to metastasis [4–6]. These hybrid E/M states facilitate collective migration [7, 8], and can confer enhanced capabilities such as stemness, drug resistance, and immune evasion to metastatic cells [6, 9, 10]. Thus, characterization and regulation of hybrid E/M phenotype is an active area of research [9, 11]. Experiments *in-vitro* and *in-vivo* have shown that hybrid E/M cells have a lower abundance in a heterogeneous population of cells, and they transition to epithelial or mesenchymal phenotypes with relative ease when compared to epithelial or mesenchymal cells [6, 12]. This weaker stability is a double-edged sword, making hybrid cells highly adaptable to metastatic stress, and also making it harder to capture and characterize these states experimentally. Advancements in high-throughput single-cell measurements have made it possible to capture hybrid E/M cells in high resolution. The two consistent characteristics of hybrid stats observed in experiments are as follows: 1) hybrid E/M cells express both epithelial and mesenchymal genes. 2) The expression levels of epithelial (mesenchymal) genes in hybrid E/M cells are lower than those in epithelial (mesenchymal) cells; in other words, genes in hybrid E/M cells are partially expressed [5, 6, 9]. Due to their limited abundance, a mechanistic understanding of the origin and expression patterns of hybrid E/M phenotypes is still lacking.

Gene regulatory networks (GRNs), defined as networks of transcriptional and post-transcriptional regulation of transcription factors and microRNAs, play a crucial role in orchestrating the cell-fate transitions underlying EMP [13]. Mathematical models of GRNs have therefore been instrumental in elucidating tumor progression and the emergence of diverse phenotypes, including hybrid E/M states [14–16]. Furthermore, these models simulate the dynamics and steady-state behaviors of genes, with steady states representing experimentally observed phenotypes. GRN models are broadly classified as continuous or discrete, based on the allowed gene expression levels. Continuous models, typically based on ordinary differential equations (ODEs), describe the production and degradation kinetics, representing gene expression as continuous positive values. Discrete models, such as Boolean formalisms, restrict gene expression to a limited number of discrete levels (high or low in Boolean formalisms). Continuous models, typically based on ordinary differential equations (ODEs), offer greater accuracy in capturing the dynamics of gene expression but require detailed knowledge of kinetic parameters, which are often unavailable for complex networks [17]. In contrast, Boolean models are simpler and rely solely on network topology, that is, the nature and direction of the regulatory interactions. Their binary framework, which assumes genes are either fully active or inactive, is supported by the sigmoidal, switch-like behavior of biomolecular interactions, such as transcription factor binding to DNA [17]. Consequently, smaller phenomenological networks of EMP have employed ODE-based models to simulate detailed dynamics [18, 19], while larger networks, which capture greater regulatory complexity, have utilized Boolean formalisms [14, 20–23].

Continuous ODE-based models of EMP networks demonstrate stable hybrid phenotypes with partial expression of epithelial and mesenchymal genes [19, 24], as observed experimentally. However, traditional Boolean models, limited to binary expression levels, cannot capture this partial expression. Instead, they identify hybrid states characterized by high expression of subsets of epithelial and mesenchymal genes [14, 15, 21]. Furthermore, the hybrid states captured by Boolean models of EMP are much less stable, as they tend to lose their phenotype quickly in the presence of external noise [15, 21]. However, hybrid E/M cells undergoing metastasis must maintain a certain level of robustness, in addition to being adaptable to environmental stress. Collective migration, which requires cancer cells to both adhere and migrate, would not be possible if hybrid states readily transitioned to fully epithelial (E) or mesenchymal (M) phenotypes, as observed in Boolean models. To address this limitation while preserving the simplicity of Boolean approaches, here we present a multilevel Boolean formalism that incorporates intermediate gene expression levels. Our simulations demonstrate that this formalism generates a greater number of steady states than the traditional Boolean model, crucially capturing hybrid E/M states with partial expression of both epithelial and mesenchymal genes. Furthermore, by introducing a modified simulation approach, we show that these hybrid states are more robust than those identified in traditional Boolean models.

## 2 Results

### 2.1 Partial expression of genes in transcriptomic data

The existence of hybrid E/M states (or simply hybrid states) was first proposed in studies of small networks modeled as sets of coupled ODEs [19, 26], and later observed in various cell lines [27, 28] and patient data [29, 30]. Furthermore, these hybrid states were found to be remarkably stable, persisting for extended periods in laboratory experiments [27], despite being relatively rare in heterogeneous EMP populations [28]. Hybrid cells are particularly relevant in the context of metastatic disease, as their plasticity facilitates secondary tumor growth and drug resistance [31]. From these studies, the basic profile of a hybrid E/M state emerged: when subjected to EMP induction, a fraction of the population retained intermediate levels of epithelial markers while also expressing intermediate levels of mesenchymal markers. Subsequently, these properties were observed in larger networks across a large parameter space [24].

Similar partial expression of epithelial and mesenchymal markers is observed in transcriptomic data. As an example, **Figure 1A** shows the expression levels of EMP genes [25] in carcinoma cell lines obtained from the CCLE (Cancer Cell line expression) database [32]. Each row corresponds to the bulk RNA-seq expression levels of a cell line, and each column corresponds to a gene, labelled on the x-axis. We classified each cell line into epithelial, mesenchymal, and hybrid phenotypes using scores calculated from the expression matrix (**Figure 1B**, Methods) and found a higher abundance of E and M samples than hybrid samples. The hybrid cell lines also show a coexpression of epithelial and mesenchymal genes, whereas the epithelial cell lines express predominantly epithelial genes and the mesenchymal cell lines express predominantly mesenchymal genes. Bulk RNA seq data showing co-expression of E and M genes is primarily composed of single cells co-expressing E and M genes [27]. We further investigated the expression levels of each gene by defining intermediate levels of expression to range from 25^*th*^ to 75^*th*^ percentile of their corresponding expression vectors across all samples. Notably, in more than 80% of the hybrid cell lines, at least 50% of epithelial and mesenchymal genes showed partial expression (**Figure 1C**). We also tested other ranges to label the expression of a gene as partial, and found that in all those ranges, the extent of partial expression was higher in hybrid samples than in epithelial and mesenchymal samples (**Figure S1**).

**Figure 1:**
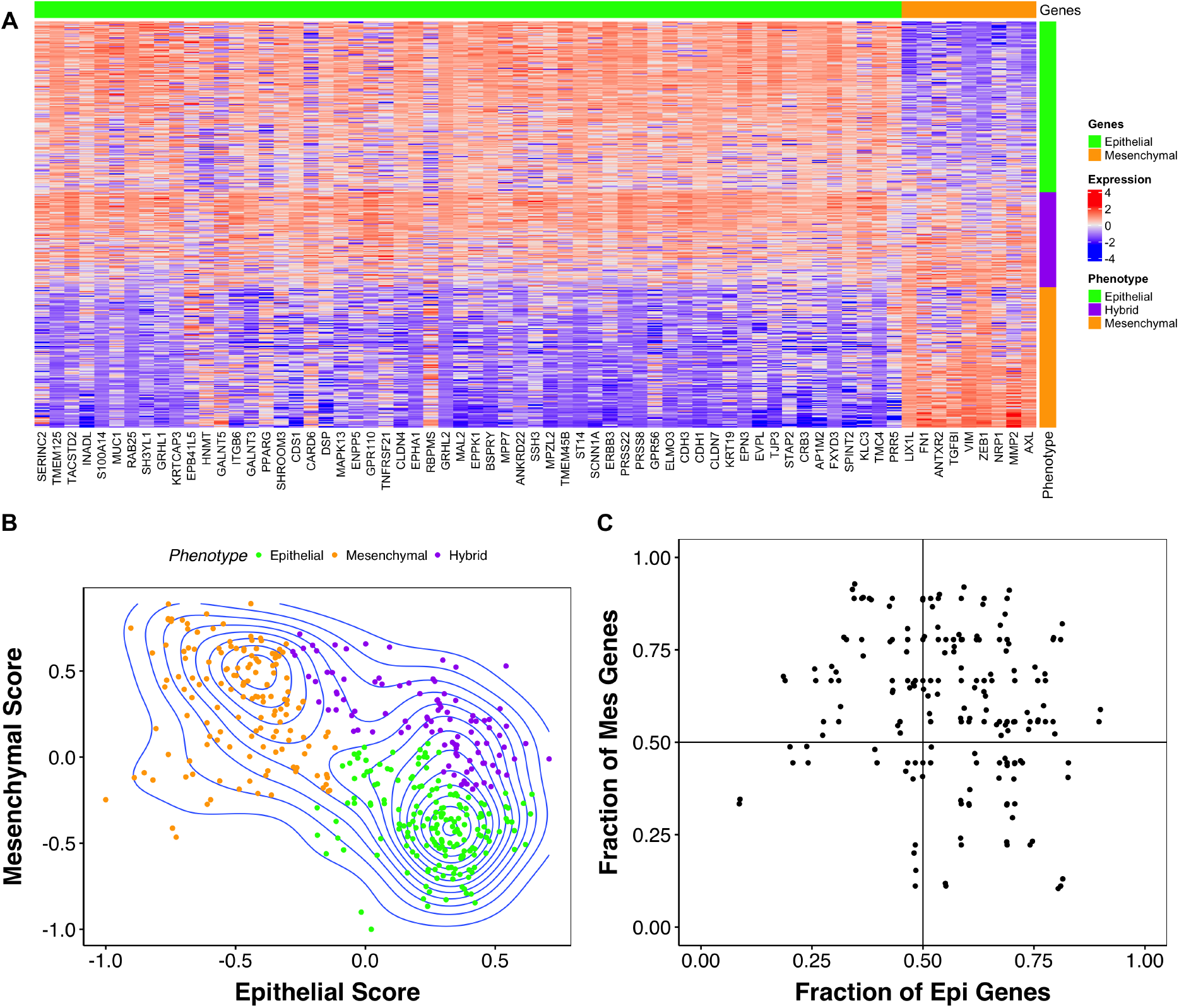
Partial expression of genes in CCLE RNA seq data. **A)** Log-normalized expression levels of CCLE carcinoma cell lines for an EMP geneset [25]. **B)** Scatterplot depicting the epithelial and mesenchymal scores calculated from the expression matrix in **A)**. Each point is a sample colored by its corresponding phenotypic classification. The contours represent the density of points at each coordinate, with smaller circles indicating higher density. **C)** Scatterplot showing the fraction of epithelial and mesenchymal genes with intermediate expression levels (between 25th and 75th quantile). Each point is a sample characterized as being in a hybrid state.

However, Boolean models are unable to capture hybrid states with partial expression of E and M genes. The framework Booleanizes the expression levels of the nodes, and as a consequence, hybrid steady states captured by Boolean models fully express subsets of both epithelial and mesenchymal genes and fully suppress other epithelial and mesenchymal genes. Hence, we asked whether the classic Boolean formalisms can be extended to capture the partial expression of E and M genes in hybrid phenotypes.

### 2.2 Multilevel models capture stable hybrid states with partial expression of epithelial and mesenchymal nodes

A Classic threshold-based Boolean formalism assumes the following dynamics :

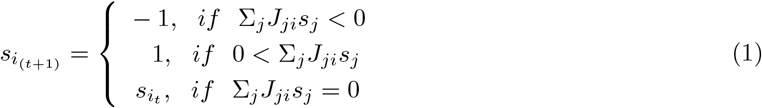

Where *s*_*i*_(*t*) is the expression level of the *i*^*th*^ gene/node (please note that we use “gene” and “node” interchangeably from here on) in the network that can take two values: -1 and 1. *J*_*ji*_ is the weight of the regulatory interaction (referred to as an edge) from the *j*^*th*^ node to the *i*^*th*^ node and takes the value -1 for inhibiting edge, 1 for activating edge, and 0 for no edge. Various models of this form, with different network structures (i.e., the set of *J*_*ji*_’s), have been used to study EMP [14, 21, 33, 34]. These models have successfully captured the properties of the two dominant phenotypes (E and M states) and the transitions that occur, such as those seen when certain state-specific factors are overexpressed.

As the Boolean formalism in equation 1 only allows two levels of expressions that represent on (+1) or off (-1), we started by introducing additional levels of allowed expression. Our specific choice was meant to maintain the symmetry of the expression levels, and therefore we introduced two new levels: -0.5 and 0.5, such that each node can take four values: −1, −0.5, 0.5, *and* 1. We define the corresponding update rules as follows:

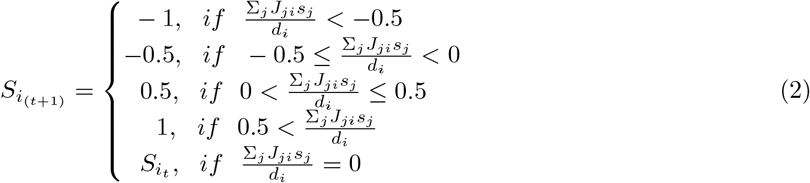

Here, *d*_*i*_ is the in-degree of *i*^*th*^ node. From here on, we refer to the model defined by Equation 1 as the two-level model, allowing two levels of expression: -1 and 1. Similarly, we refer to the model defined by Equation 2 as the four-level model. To characterize the effect of the four-level model on the steady states, we applied the two-level and four-level models to five different EMP networks from the literature [14, 21–24, 35]. These networks have a range of sizes (smallest at 20 nodes (N) and 40 edges (E), and largest at 72N 142E) and, more importantly, are constructed in varying biological contexts. The first pair of networks - 22N 82E and 26N 100E were chosen to analyze the effect of phenotype stability factors (PSFs): OVOL2 and NP63, which have been shown experimentally and through mathematical models to stabilize hybrid E/M phenotypes [27]. The 20N 40E network was constructed in the context of Breast cancer proposing additional PSFs [22], 31N 95E network in the context of Lung cancer studying the dynamics of EMT reversal [23], and the 72N 142E network was developed in the context of hepatocellular carcinoma [14] to study TGF*β* driven Epithelial Mesenchymal Transition (EMT) and dysregulated signalling pathways in HCC, which was further extended to include pathways of reversal of EMT [21]. Using these networks, we aimed to analyze the emergent hybrid states in the presence of varying topological as well as biological contexts.

We focus primarily on the 26N 100E EMT network [35] (**Figure 2A, top**). EMT networks can be visualized as having two mutually inhibiting groups or “teams” of nodes. One team consists of epithelial nodes (micro-RNAs, CDH1, CLDN7, GRHL2), and the other consists of mesenchymal nodes (FOXC2, GSC, KLF8, SNAI1, SNAI2, TCF3, TGFbeta, Twist1, Twist2, Zeb1, and Zeb2) respectively. As the nodes within the same team belong to the same phenotype, the interactions within a team (that is, epithelial to epithelial and mesenchymal to mesenchymal) are predominantly positive, while those between teams (epithelial to mesenchymal and mesenchymal to epithelial) are predominantly negative, thereby leading to a higher-order toggle switch between the two teams (**Figure 2A, bottom**). This structure strongly encourages the coexpression of nodes in the same team and antagonistic expression patterns between the two teams. Consequently, the team structure reduces the probability of the occurrence of hybrid states, which require the coexpression of nodes from both E and M teams [15]. In addition to the teams, this network has genes known as PSFs (phenotype stability factors [27]): NP63 and Ovol2, which do not align with any team and therefore can stabilize hybrid phenotypes. Due to these features, this network can indeed exhibit stable, low-frequency, hybrid phenotypes. For ease of analysis, we removed the input nodes that do not have incoming edges from any other nodes in the network, and the output nodes that do not have any outgoing edges in the network (gray colored nodes in (**Figure 2A**). Output nodes (VIM in the 26N 100E network) do not affect any other nodes in the network and hence do not have any effect on the dynamics. Input nodes create inconsequential degeneracy in the steady states. For example, simulating the 26N 100E network using the two-level model results in four unique configurations of epithelial phenotype and four unique configurations of mesenchymal phenotype (**Figure 2B**). However, the only difference within the four epithelial states (and within the four mesenchymal states) is the difference in expression of the signal nodes, highlighted in a black box. Each epithelial (mesenchymal) configuration has the same abundance. Thus, after removing the input and output nodes, the resultant network has 23 nodes and 90 edges (23N 89E). We eliminated the input and output nodes from the remaining four networks: the network without the PSFs (22N 82E, 15N 60E without the peripheral nodes, **Figure S2A**), the breast cancer EMP network [22] (20N 40E, 16N 32E without the peripheral nodes, **Figure S2B**), the lung cancer EMP network [23] (31N 95E, 21N 77E without peripheral nodes, **Figure S2C**) and the large HCC EMP network [21] (72N 142E, 55N 111E without peripheral nodes, **Figure S2E**). The edge tables corresponding to these networks are given in **Supplementary Table 1**.

**Figure 2:**
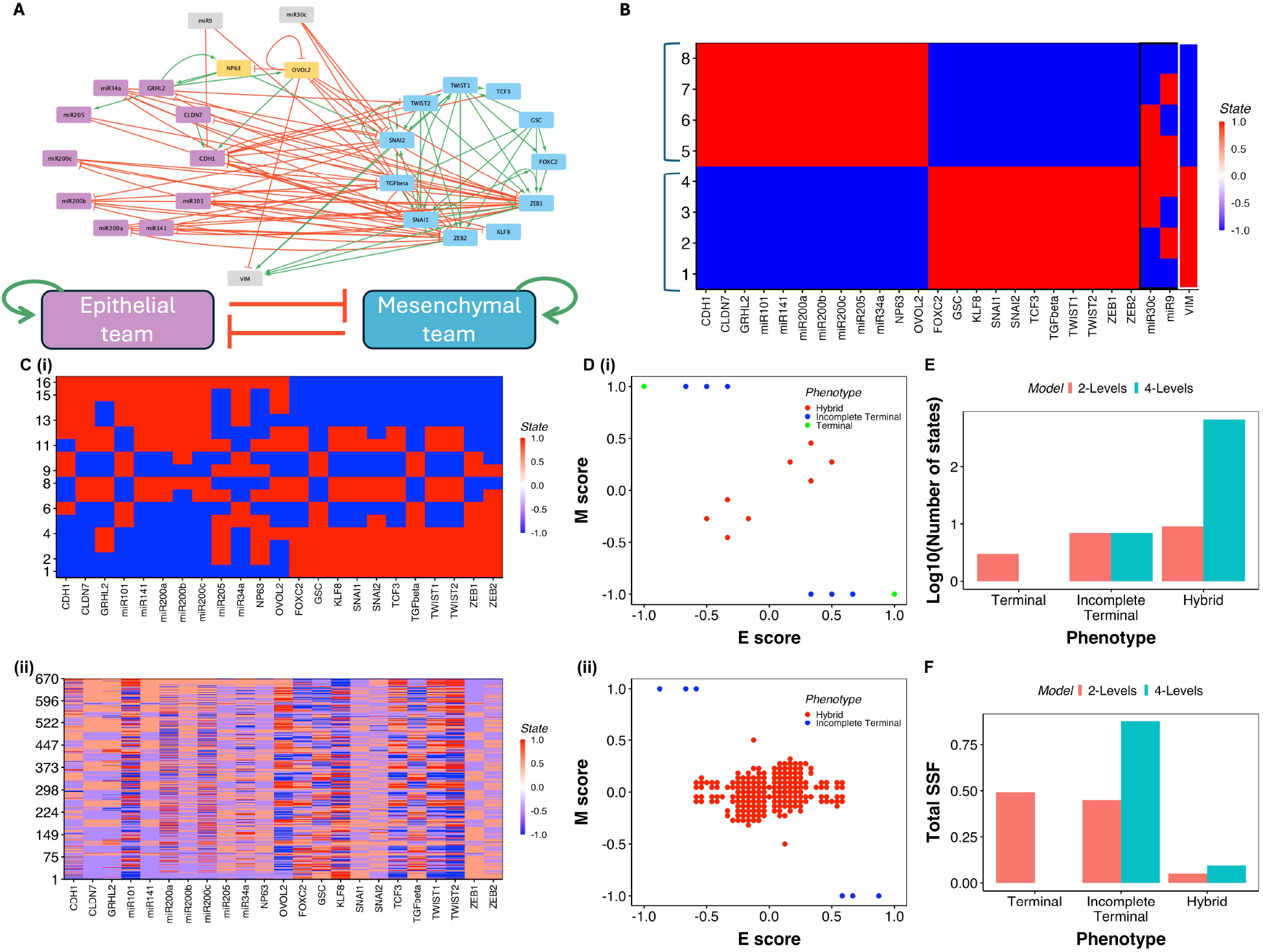
Multilevel formalism results in an increased number of unique hybrid states. **A)** The 26N 100E EMP network, and the schematic of the team abstraction of the network. The pink nodes are epithelial, the blue nodes are mesenchymal, the gray nodes are peripheral nodes, and the yellow nodes are the PSFs. **B)** Configuration of epithelial and mesenchymal states obtained by simulating the 26N 100E network with a two-level model. Input nodes and their expression are highlighted with a black box, and the output node is highlighted with a white box. **C)** Heatmaps depicting the configurations of steady states of the 23N 89E EMP network simulated using i) two-level and ii) four-level threshold-based formalism. Each row corresponds to one steady state, and each column corresponds to a node labeled at the bottom. Red corresponds to a positive expression level, and blue corresponds to a negative expression level. **D)** 2-D scatter plots of the steady states obtained using i) two-level and ii) four-level threshold-based formalism, on the E and M score axes. Each point is a steady state. **E)** Barplot depicting the comparison of the number of steady states (y axis) of each category (hybrid, Terminal, and Incomplete terminal, x-axis) between two-level formalism (red) and four-level formalism (green). **F)** Same as C, but for the total frequency of all steady states of a category.

Using Equations 1 and 2, we simulated each network starting from 10^7^ random initial conditions (RIC) till each RIC converged to a steady state. Each RIC represents a cell in a heterogeneous population, with the corresponding steady state being analogous to the phenotype of that cell (**Figure 2A**). This ensemble of steady states is analogous to the transcriptomic dataset presented in **Figure 1A** (compare with **Figure S4A**). From the ensemble of 10^7^ steady states (**Figure S4A**), we identified the unique steady states (16 in the two-level model and 670 in the four-level model from 10^7^ RICs) and their frequencies for further analysis. To classify these states/phenotypes, we define epithelial and mesenchymal scores as the average expression of epithelial and mesenchymal nodes, respectively, for a given steady state (see **methods** and **Supplementary Table 2** for the list of epithelial and mesenchymal genes for the different EMP networks). We then defined a phenotypic score as the difference between the epithelial and mesenchymal scores for a given steady state. While the epithelial and mesenchymal scores range between *−*1 and 1, the phenotypic score ranges between *−*2 and 2. The epithelial (mesenchymal) states, defined as the states with the high expression level (1) of only epithelial (mesenchymal) nodes, and low expression level (*−*1) of all mesenchymal (epithelial nodes, have +2(*−*2) phenotypic score. Similar to the transcriptomic data, our simulations result in a high abundance of epithelial and mesenchymal states in the top-left and bottom-right corners of the epithelial-mesenchymal score space (compare **Figure 1B** with **Figure S4B**). We classified the emergent steady states into three classes based on the phenotypic scores, namely, terminal, incomplete terminal, and hybrid. Terminal steady states are the states with phenotypic score +2 or *−*2, which include the pure epithelial and pure mesenchymal states described earlier. Note that these states have symmetrical composition, that is, nodes that are turned off in epithelial states are turned on in mesenchymal states and vice versa. Incomplete terminal states are the steady states with either epithelial score or the mesenchymal score equal to *−*1, and the corresponding mesenchymal or epithelial score less than +1. In other terms, these are the steady states with incomplete expression levels of either only epithelial or only mesenchymal nodes. We also include the symmetrical case, where the epithelial or mesenchymal score is +1 and the corresponding mesenchymal or epithelial score is negative. Although these states exhibit co-expression of epithelial and mesenchymal genes, since all nodes from one team are completely active, the properties of the corresponding phenotype dominate [6]. Our distinction of terminal and incomplete terminal states is aimed at capturing the heterogeneity in the expression patterns of experimentally observed epithelial and mesenchymal phenotypes. Biologically, both terminal and incomplete terminal phenotypes have similar functions: epithelial and incomplete epithelial phenotypes are expected to be strongly adhesive, while mesenchymal and incomplete mesenchymal phenotypes are highly migratory. The remaining steady states that could not be classified as Terminal or Incomplete terminal express subsets of both epithelial and mesenchymal nodes, and were classified as “hybrid” steady states. Note that the hybrid states have a low magnitude of phenotypic scores as compared to Terminal and Incomplete terminal states (**Figure 2B**).

The four-level model results in a larger number of unique steady states (670 in the four-level model versus 16 in the two-level model, **Figure 2C**). While the number of terminal and incomplete terminal states together in the two-level model was comparable to that of hybrid states (5 vs 11), the four-level model results in a much larger number of hybrid states (compare the centers of **Figure 2D, i and ii**). Of the 670 steady states observed in the four-level model, 664 are hybrid states, a 66 fold increase from the two-level model(**Figure 2E**). Note that these hybrid states have intermediate levels of epithelial and mesenchymal scores, indicating coexpression of subsets of epithelial and mesenchymal genes as observed experimentally. The increase in the number of steady states does not scale with the state space, even for higher number of levels (2^2^3 to 4^2^3, **Figure S7A**), suggesting that the newly appeared hybrid states may not just be an artifact of the increase in the state space size. We further simulated a randomized version of the 23N 89E network, generated by shuffling the signs of the edges in the network (thereby maintaining all properties of connectivity of the network), and we observed more than 2100 steady states from four-level formalism, as compared to 32 steady states in the two-level formalism (**Figure S4**). On the other hand, the network obtained by removing PSFs from the 23N 89E network (15N 60E) showed only 90 steady states in the four-level formalism (**Figure S5A**). Thus, the number of steady states resultant from four-level formalism remains strongly dependent on the structural and functional features of the network being simulated.

We next investigated the abundance of the hybrid and non-hybrid phenotypes in the ensembles generated from the simulations. We calculated the sum of the steady-state frequency (SSF) for all hybrid states, the indicator of the abundance of the hybrid phenotype in the digital cell population. Although the number of unique hybrid steady states showed a 66-fold increase from the two-level to the four-level model for the 23N 89E network, the total abundance of hybrid phenotype showed only a two-fold increase from 0.05 to 0.1. Similar observations were made for the four other EMP networks analyzed here (**Figure S5, S6**). In the 15N 60E network (23N 89E network without the PSFs), only two hybrid states with negligible frequency were found using the two-level model. The number of hybrid states increased to 90 using the four-level formalism (i.e., 45 fold change), and the total frequency of hybrid states increased from 0.05 to 0.3 (**Figure S5A**). In the 16N 32E network, we observed a 10-fold increase in the number of hybrid states (12 to 124) while the corresponding frequency increased from 0.4 to 0.7 (**Figure S5B**). In 27N 32E and 55N 111E networks, we found a much larger increase in the hybrid state frequency (**Figure S6**). The frequency of terminal and incomplete terminal states decreased and the abundance of hybrid states increased with increasing fraction of weighted negative loops (FWNC), a measure of the extent of negative feedback in a network, calculated as 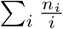, where *n*_*i*_ is the number of negative loops with *i* edges [16, 36] (**Figure S8**, Supplementary methods), highlighting the topological origins of hybrid states. Across the EMP networks, we found that the number of unique hybrid states from the two to four-level model increases disproportionately to the total abundance of hybrid states. These observations indicate significant heterogeneity in the stability of the newly obtained hybrid states and suggest that, in general, these states individually are much less stable than terminal and partial terminal states that show a higher abundance with a smaller number of unique states.

### 2.3 Frustration and steady state frequency (SSF) of four-level hybrid states indicate improved stability

We further probed the stability of the steady states using the SSF and the concept of frustration. SSF of a steady state is calculated as the fraction of the 10^7^ initial conditions that evolve into the steady state, with higher SSF indicating higher stability of the steady state. Frustration measures the extent of dissatisfaction in the network structure for a given steady state. Each edge in the network models has one of three weights: *−*1, 1, *or* 0, and a corresponding “expectation” for the expression levels of the source and target nodes - opposite expression levels for an inhibitory edge (weight of *−*1) and similar expression levels for an activating edge (weight of +1). An edge is frustrated if these expectations are not met. Specifically, if the product of its weight and the expression levels of the nodes participating in the edge is negative (i.e., *J*_*ji*_*s*_*i*_*s*_*j*_ *<* 0)(**Figure 3A, i**). We define the frustration of a steady state as the sum of frustration of all edges in the network divided by the total number of edges 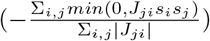 (equation 10). Frustration correlates negatively with the stability of a steady state [15, 34]. As the dynamics (update rules, equations, or parameters) align with the network structure, one can intuitively infer that lesser conflict of a state configuration with the edges in the network would lead to higher stability. Specifically for the two-level formalism, we can show that steady states cannot have a frustration beyond 0.5. From equation 1, the stability condition for a node can be derived as 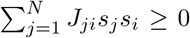, which is only achievable if less than half the incoming edges are frustrated. Consequently, perturbations targeting frustrated edges have a higher chance of transitioning to a new state, as those transitions can increase the sum 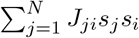, thereby enhancing the stability of the node and the state (see the example in **Figure 3A, ii**). In the two-level model of the 23N 89E network, the states with a high magnitude of the phenotypic score (*>* 1) show low frustration, and vice-versa (**Figure 3A, i**). The four-level model leads to a decrease in the overall frustration of the steady states in all ranges of the phenotypic score (**Figure 3A, ii**). The states with high phenotypic scores, i.e., the terminal and incomplete terminal states, maintain low frustration. Although most of the hybrid states (with low phenotypic scores, colored red) show high frustration, we find some hybrid states in the four-level model with low phenotypic scores but lower relative frustration (0.04 against a maximum frustration of 0.2) as compared to the two-level formalism (0.25 against a maximum of 0.4, **Figure 3A, i vs ii**). Thus, the four-level formalism allows for a larger number of low-frustration hybrid states, signalling an increase in their stability. The frustration distributions of the terminal and incomplete terminal states are closer to that of hybrid states as compared to the two-level states, a property also observed in the CCLE data (**Figure S11**).

**Figure 3:**
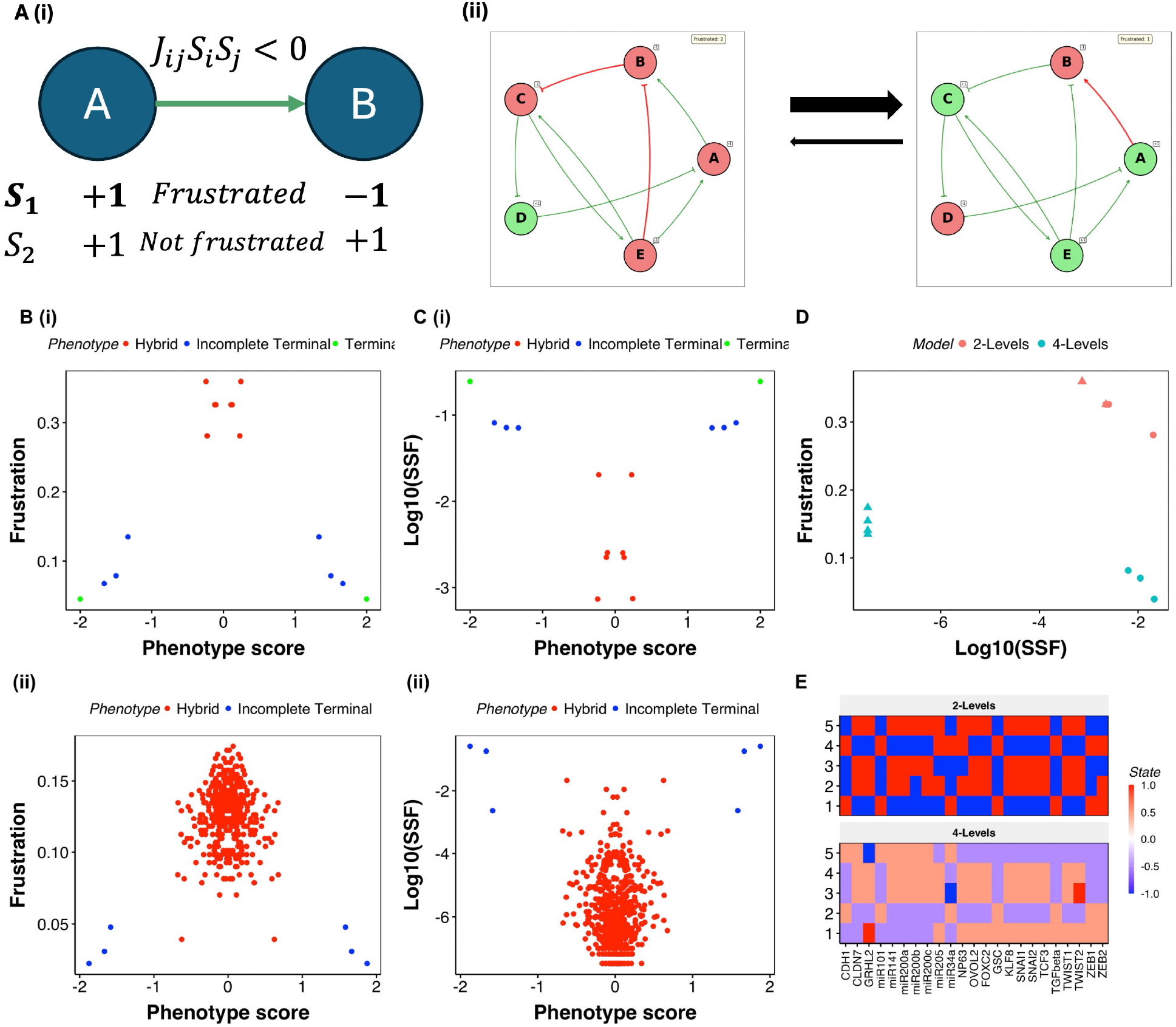
Four-level model results in similar SSF but lower frustration hybrid states with partial expression of nodes. **A)** (i) Schematic depicting the criteria for an edge to be frustrated. The activation between A and B is frustrated in a state if A and B have opposite levels of expression in that state. (ii) An example of the effect of frustration on the transition between two steady states in a toy network. Each node with expression +1 is colored green, and each with expression -1 is colored red. Activating edges end with arrows while inhibiting edges end with hammers. Edges colored in red are frustrated. **B)** Scatter plots depicting the Frustration of the steady states of the 23N 89E network simulated using i) two-level and ii) four-level formalism. **C)** Same as A, but for steady state frequency (SSF). **D)** Frustration and SSF of most frequent (circles) and least frequent (triangles) hybrid states from two-level (red) and four-level models (green). **E)** Heatmaps depicting the composition of most frequent hybrid states from two-level (top) and four-level (bottom) formalisms.

Following the frustration trends, the terminal and incomplete terminal states have a higher SSF than hybrid states (**Figure 3B**). A large fraction of hybrid states have very low SSF; however, some of the hybrid states with relatively higher frequency are found in both two-level and four-level formalisms, with both low (*<* 1) and high (*>* 1) magnitude of phenotypic scores. However, while the hybrid states of the level model have strictly lower SSF than the terminal and incomplete terminal states, a fraction of the hybrid states of the four-level model have lower but comparable SSF to that of terminal and incomplete terminal states, which further suggests that the hybrid states of the four-level model are more stable than the two-level hybrid states. Note that the simulations of EMP networks at a larger number of levels fail to capture the stability differences between hybrid and terminal phenotypes, which serves as a ground truth for our models (**Figure S7**). Hence, we restrict the number of levels to four.

We then compared the frustration and stability of the high-frequency hybrid states obtained from two-level and four-level models. The high-frequency hybrid states had lower frustration in the four-level model, and had a comparable, albeit slightly higher SSF as that of the two-level model (**Figure 3C**). The composition of these states revealed further interesting results. The hybrid states of the two-level model could be classified into two groups: states where a large number of nodes are on and those where a large number are off (states 2,3,5 vs 1,2 in **Figure 3D**, top). In the four-level model, we find all the top hybrid states with multiple E and M nodes having intermediate expression levels (-0.5, 0.5), a partial expression feature observed in hybrid samples in experimental data. Similarly, we found hybrid states with lower frustration, higher frequency, and partial expression of epithelial and mesenchymal nodes in 15N 60E, 16N 32E, and 27N 76E networks (**Figure S9, S10**). While the multilevel formalism did result in hybrid states with low frustration and high frequency in the 55N 111E network, the extent of partial expression of nodes in the top hybrid states is not as high as seen in the other networks. Overall, we find that multilevel formalism gives rise to a new type of stable hybrid state, with a partial expression of epithelial and mesenchymal nodes that are not seen in two-level formalism.

### 2.4 Multilevel hybrid states are resilient to the loss of uncertainty in simulation formalism

Our results so far have shown that increasing the granularity of the state space by employing a multilevel formalism increases the incidence of low-frustration, high-SSF hybrid states with partial expression. Previous Boolean simulations of EMP networks have shown that the network structure, involving “teams” of E and M nodes, does not tend to favor hybrid states [15], which is the primary reason for their low stability and high plasticity.

However, hybrid states can emerge through a different mechanism: regulatory uncertainty in the update rules. In biological networks, genes may sometimes lack clear regulatory signals due to balanced opposing inputs or weak regulatory contexts. The threshold-based formalism captures this through the rule *s*_*i*_(*t* + 1) = *s*_*i*_(*t*) when 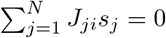.

An example of such behavior is the degeneracy in steady states introduced by input nodes in the network. As input nodes are not regulated by any other nodes, they constantly maintain their expression irrespective of the expression of downstream genes, causing degeneracy and frustration in steady states (**Figure 2B**). More generally, any node can experience this regulatory uncertainty during network dynamics. Consider a mutual inhibition loop between nodes A and B with an inhibitory signal from node C to A and a self-activation on B. When C is inactive and B is active, the net regulation on A is zero (*J*_*BA*_*s*_*B*_ + *J*_*CA*_*s*_*C*_ = *−*1 + (+1) = 0). If node A was initially active (*s*_*A*_ = +1), the net regulation on node B also is zero *J*_*BB*_*s*_*B*_ + *J*_*AB*_*s*_*A*_ = +1 + (*−*1) = 0). Thus we obtain a hybrid state, with both A and B active and the edges between A and B frustrated.

Such regulatory uncertainty allows nodes to maintain expression states that are decoupled from the current network context, and can stabilize otherwise unstable hybrid configurations by preventing the system from fully committing to either epithelial or mesenchymal states. We asked whether the stability of hybrid states in two-level and multilevel formalisms depends on the degeneracy introduced by these unregulated nodes during network dynamics. We therefore considered a modified version of the threshold-based Boolean formalism [37]:

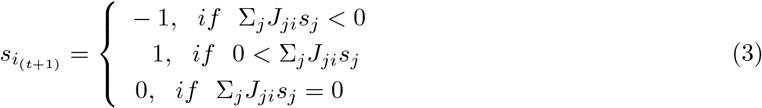

This formalism, referred to as the “turn-off” formalism from here on, sets the expression of unregulated nodes at each time step to zero, making them inconsequential in the next time step of the simulation. Therefore, unregulated nodes are given a fixed state, eliminating their ability to introduce noise in the dynamics. One immediate implication of this formalism is that the expression of input nodes (nodes with zero in-degree) is always set to zero. Biologically, zero expression level could be interpreted as a basal expression level that has no regulatory impact on the downstream nodes. As input nodes introduce degeneracy in epithelial and mesenchymal states (thereby creating microstates in the landscape that have the same level of stability), this formalism is expected to eliminate this degeneracy. To verify this hypothesis, we simulated the 26N 100E network (23N 89E network in the previous figures, plus input and output nodes) with a normal threshold and the turn-off formalism. In the two-level formalism, the 26N 100E network gives rise to 114 unique steady states, of which 8 are degenerate terminal states that have similar SSF and frustration but differ in the configuration of the input node expressions (**Figure 4A**). Indeed, we see that applying the turn-off condition selectively to the input nodes of the 26N 100E network reduces the number of steady states from 114 to 20 (**Figure 4A, B**), with a single epithelial and a single mesenchymal state (as opposed to 4 states each without node turn-off). We further extended the turn-off restriction to all nodes and found that the number of steady states decreased to four (**Figure 4C**). Among these four states, the epithelial and mesenchymal states completely take over the state space with a total frequency of 0.999.

**Figure 4:**
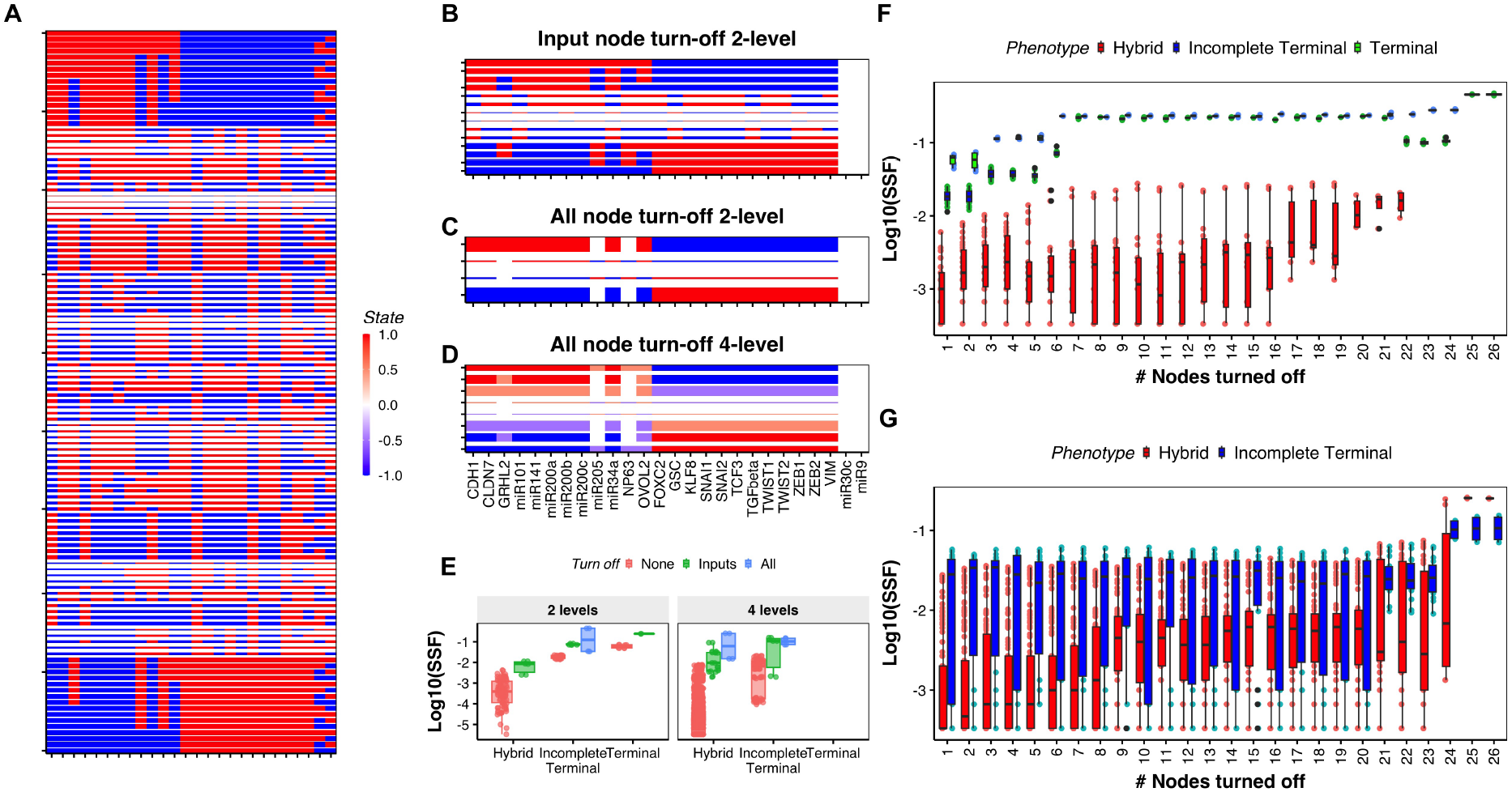
Node turn-off highlights the stability of hybrid states in multilevel formalism. **A)** Heatmap depicting the configuration of steady states of the 26N 100E network (23N 89E + input and output nodes), simulated with the two-level formalism. Red corresponds to negative expression, and blue represents positive expression. The height of each row is proportional to the log10(SSF) of that steady state. **B)** Steady states obtained by applying turn-off to input nodes with the two-level formalism. The color scale is the same as that of A. The height of each row is proportional to the log10(SSF) of that steady state. **C)** Steady states obtained by applying turn-off to all nodes with the two-level formalism. The color scale is the same as that of A. The height of each row is proportional to the log10(SSF) of that steady state. **D)** Steady states obtained by applying turn-off to all nodes with the four-level formalism. The color scale is the same as that of A. The height of each row is proportional to the log10(SSF) of that steady state. **E)** Boxplots depicting the SSF of steady states of different classes from four-level and two-level models - without turn-off (red), turn-off applied to input nodes (green), and turn-off applied to all nodes (blue). **F)** Boxplot depicting the evolution of steady state frequency distribution for the sequential turn-off experiment to optimally increase total frequency of hybrid states, simulated using the two-level formalism. **G)** Same as **F)**, but turn-off is applied to the four-level formalism.

Thus, we can conclude that the hybrid states obtained in the two-level formalism largely emerge due to the degeneracy in the dynamics. Nodes like NP63, which have conflicting interactions with the “team” structure of the network (activate both an epithelial (GRHL2) and mesenchymal (SNAI2) gene; inhibited by the epithelial gene OVOL2 but activated by GRHL2), are often the cause of the increase in degeneracy and hybridness in the steady states. We find that the turn-off formalism leads to the NP63 node being turned off in the dominant steady states of the 26N 100E EMP network.

The turn-off formalism also reduces the number of states that emerge from the four-level formalism to 8 (**Figure 4D**). Among these, the first two and the last two are incomplete terminal states, while the ones at the center are hybrid states. As most of the states emerging from the multilevel formalism are weakly stable hybrid states, the turn-off formalism readily eliminates these states. However, the most dominant hybrid states see a significant increase in their frequencies (from the order of 10^*−*3^ without turn-off to 10^*−*1^ with the turn-off condition applied to all the nodes - **Figure 4E**). These hybrid states are composed of partial expressions (-0.5 or 0.5) from nearly all the nodes in the network; the states that we have previously referred to as “true” hybrid states. While the true hybrid states were lagging in SSF and frustration compared to other hybrid states with limited partial expression in the absence of turn-off (**Figure 3A ii, B ii, and D**), turn-off formalism leads to enhanced stability of the true hybrid states, asserting their significance. Thus, by reducing the degeneracy allowed in the simulation, the turn-off formalism can greatly concentrate the steady states of a network. Adding the relaxation of multilevel formalism to the state space allows for the concentration of hybrid states, establishing the stability of these “true” hybrid states that cannot be captured in the two-level formalism.

As we have just shown, applying turn-off to input nodes alone retained multiple weakly stable states, while applying turn-off to all nodes eliminated most of the weakly stable states in both two and four-level models. But we also noticed that nodes like NP63 were turned off in all steady states, despite not being input nodes. We were therefore interested in asking if it is possible to identify a small subset of nodes to apply turn-off to that can achieve the same results as applying turn-off to all nodes. With both the two-level and the four-level models, we carried out a “sequential-turn-off” experiment. The simulations are carried out to a maximum of N iterations (N = 26 for the 26N 100E network). At the first iteration, we apply turning off one node at a time and identify the node *N*_1_ that gives the maximum “hybridness” calculated as the total frequency of hybrid states. In the next iteration, we apply turn-off to two nodes at a time to identify the next node *N*_2_ that, when turned off along with the other, *N*_1_, gives the maximum hybridness.

Interestingly, while optimizing for hybridness, we observed an increase in the mean SSF of the terminal and incomplete terminal phenotypes in both the two-level and four-level models (**Figure 4F, G**). Another interesting observation is that while the increase in mean frequency for hybrid states in both two-level and four-level formalisms happened at a similar fraction of nodes turned off, the frequency rose sharply in the four-level formalism, while that in the two-level formalism saw a gradual rise. From 24 nodes onwards, the hybrid states - with partial expression of E and M nodes - had comparable SSF to that of the incomplete terminal state. We further simulated the 22N 82E, 20N 40E and 32N 95E networks using the turn-off formalism (**Figure S12, S13**). Despite the differences in the network sizes and structures, application of turn off to all nodes consistently highlighted the stability of hybrid states with partial expression of E and M nodes. Together, these results highlight that the hybrid states obtained with the four-level formalism are more stable than those obtained from the two-level formalism.

### 2.5 Multilevel formalism captures a resilient cloud of hybrid states

One of the key features that make hybrid states conducive to metastasis is plasticity — their ability to readily change phenotype when exposed to stress. We found that multilevel hybrid states exhibited lower frustration than two-level hybrid states. Since higher frustration can make a state more susceptible to perturbations, we next asked whether this reduction in frustration affected the plasticity of four-level hybrid states. Although the steady-state frequency (SSF) of four-level hybrid states was similar to that of two-level hybrid states, SSF is not necessarily a reliable indicator of how sensitive steady states are to perturbations. Therefore, we performed two perturbation experiments on hybrid states from both the two-level and four-level models of the EMP networks to investigate their transition dynamics.

In the first experiment, we created a digital population of 100 cells, with each cell’s state chosen uniformly from a set of 10 hybrid states — specifically, the five hybrid states with the highest SSF values and the five with the lowest. We then exposed each cell in this population to transient perturbations by randomly selecting a set of *P* nodes of size *k*; 1 *≤ k ≤ N* nodes, and generated a perturbed population *S*_*P ert*_, such that 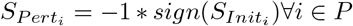 where *S*_*Init*_ represents the state configuration of a cell in the initial population. After perturbation, we allowed the population to converge to a steady state (**Figure 5A**). We repeated these experiments for 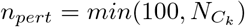 times, generating an ensemble of as many final populations. Biologically, this experiment tracks the plasticity of hybrid states when exposed to transient signals. Because the perturbed nodes are chosen randomly, the resulting perturbations can push the system toward either an epithelial or a mesenchymal phenotype. The conventional understanding of hybrid E/M state plasticity suggests that, upon perturbation, the population readily transitions to either an epithelial or a mesenchymal phenotype—labeled in our simulations as Terminal and Incomplete Terminal states [6, 12, 15]. Both models are consistent with this conventional view: the majority of the population shifts to a terminal or incomplete terminal state under moderate to large perturbations (**Figure 5B, S17A, S18A**). As expected, the maximum retention of hybridness occurs at the lowest level of perturbation, with more than half of the population remaining in hybrid states. Retention is higher in four-level hybrid states(*≈* 75%) compared to two-level hybrid states (*≈* 65%). As a control, we repeated the experiment with a population of terminal and incomplete terminal states and found that 100% of the cells retained their terminal/incomplete terminal phenotype under small perturbations (**Figure S4A**). However, in the two-level formalism, the fraction of hybrid retention drops sharply—falling below 5% in the two-level formalism once more than 50% of the nodes are perturbed. By contrast, in the four-level formalism, more than 25% of the population retains hybridness even at the maximum perturbation level.

**Figure 5:**
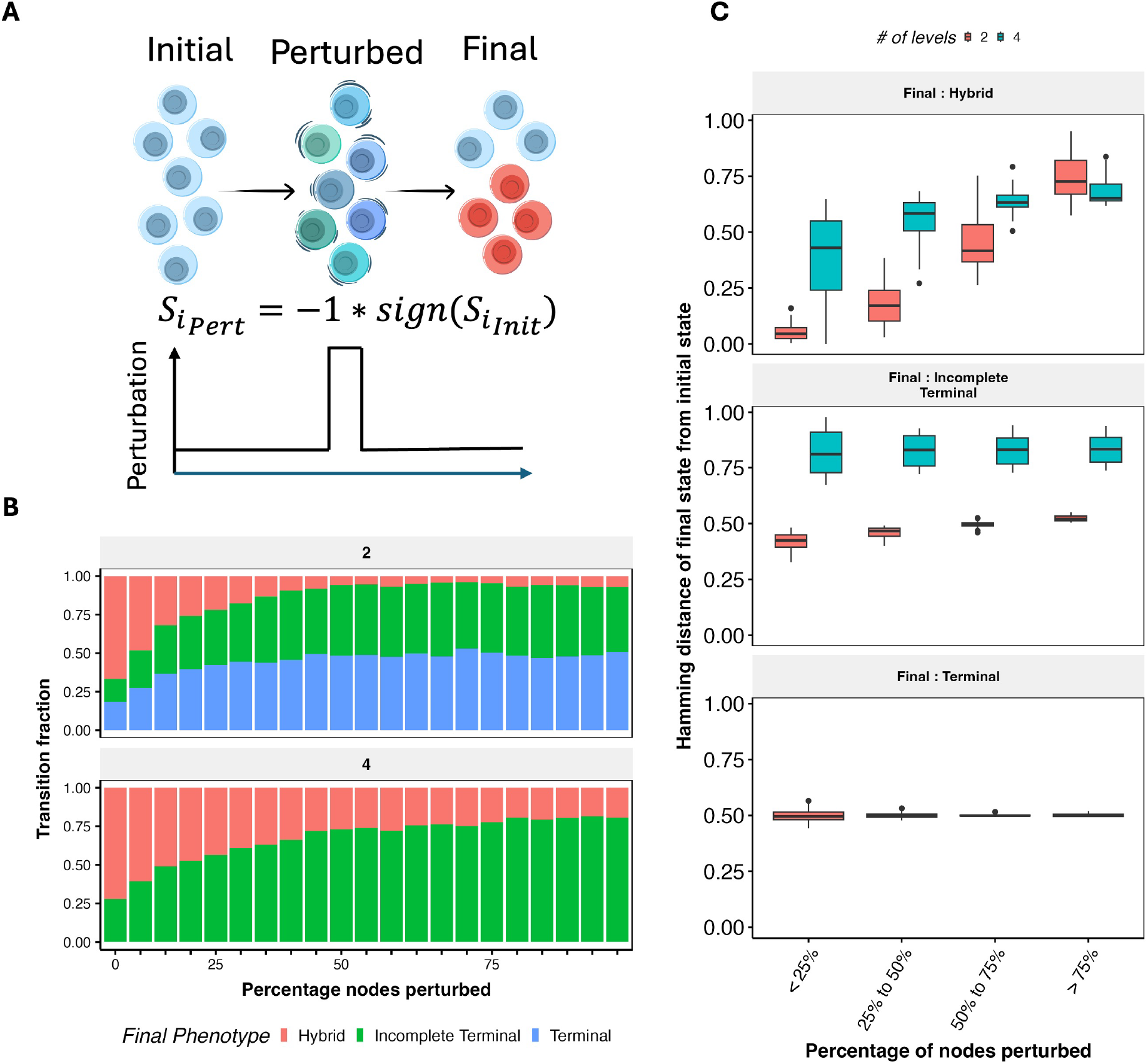
Multilevel hybrid states are more likely to retain their hybridness than two-level hybrid states. **A)** Schematic of the perturbation experiment. A population of hybrid states is exposed to a random transient perturbation, and the population is allowed to reach a steady state. **B)** Fraction of the population of hybrid states reaching Terminal, Incomplete Terminal, and Hybrid states after perturbation in 2-level (top) and four-level (bottom) models. The X-axis shows the percentage of nodes perturbed at a time, as shown in the schematic. **C)** Hamming distance of the initial hybrid states from the final state after perturbation, calculated as the number of nodes with different expression levels between the initial and final states. Each panel shows a different final phenotype achieved by the population of hybrid states.

This difference in hybridness retention between two-level and four-level models is particularly interesting because the SSFs of hybrid states—reflecting their abundance in a heterogeneous population—are similar in both two-level and four-level models. This led us to ask: how far do cells travel in expression space to reach a new steady state after perturbation? In other words, is hybrid retention in multilevel model explained by the stability of individual hybrid states, possibly due to the lower frustration of four-level hybrids, or is another mechanism at play?

To address these questions, we calculated the Hamming distance between initial and final steady states, which measures the fraction of nodes with different expression values between two states. The results for each final phenotype are shown in **Figure 5C**. Several general trends are worth noting. First, terminal and incomplete terminal states are farther away from hybrid states in the four-level model than in the two-level model, reflecting the partial expression patterns that characterize hybrid states (**Figure 5C, S17B and S18B, middle panels**). Second, the distance from hybrid to terminal/incomplete terminal states is less variable than the distance within hybrid states in both models. When the final phenotype is a hybrid, additional trends emerge. At low perturbation levels (*<* 25%), two-level hybrid states that retain a hybrid phenotype primarily do so by returning to their exact initial state, resulting in a near-zero Hamming distance. By contrast, four-level hybrids move significantly farther away even under small perturbations (mean hamming distance of 0.4, i.e., expression of 40% of the nodes changes on average). Note that at higher perturbation values, while the hamming distance in two-level model also increases, the probability of hybrid to hybrid transition is much lower than the four-level model. Consistent with the increased hamming distance, convergence times for hybrid-to-hybrid transitions are longer in the four-level formalism than in the two-level formalism (**Figure S4B, S17C, S18C**). Interestingly, transitions where both the initial and final states are terminal or incomplete terminal take less time in the four-level model than in the two-level model (**Figure S4C**). A similar observation was reported in a previous study of EMT landscapes, where the presence of hybrid states was shown to facilitate faster epithelial-to-mesenchymal transitions [38].

In summary, four-level hybrid states are more sensitive to perturbations than two-level hybrid states, making them more plastic. Yet, they retain hybridness more frequently by transitioning to other hybrid states. Moreover, the final states reached are often well separated from the initial states. Together, these results suggest that although individual hybrid states are not highly stable, the collective ensemble of hybrid states—or the “hybrid cloud”—is stable. This form of plasticity may better capture the metastatic potential of hybrid states, where enhanced plasticity promotes adaptiveness while preserving the advantageous hybrid phenotype. Encouraged by these results, we next asked how four-level hybrid states behave in a noisy environment that cancer cells would encounter during metastasis. As before, we started with a digital population of hybrid states, but this time introduced a small, recurring perturbation: every 10 time steps, one randomly selected node was perturbed in the manner described in **Figure 5A**. We then simulated the system for 10,000 time steps— *≈*100 times longer than the average convergence time to a steady state in the 23N 89E network, and *≈* 1, 000 times longer than the average convergence time to terminal and incomplete terminal states.

Interestingly, despite the low level of perturbation, two-level hybrid states quickly drifted toward epithelial or mesenchymal phenotypes (**Figure 6A (i)**, phenotypic score approaching 2 and *−* 2). To quantify this migration, we calculated the residence times of these trajectories, i.e., the number of time steps trajectories spent in different regions of the E-Score–M-Score plane. A density map of residence times across 1,000 trajectories confirmed that most trajectories localized to the corners corresponding to epithelial and mesenchymal phenotypes (**Figure 6B**).

**Figure 6:**
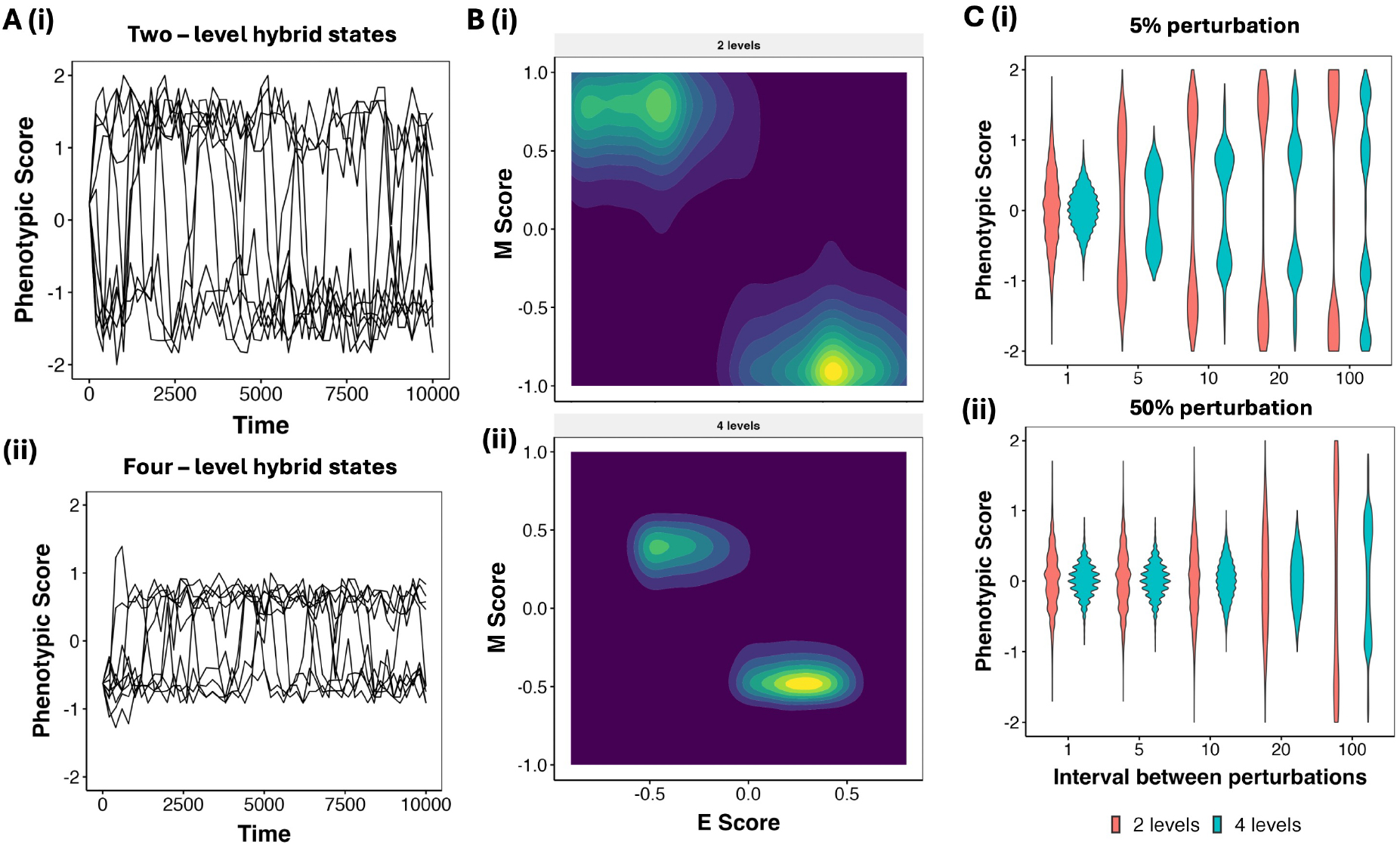
Resilience of the multilevel hybrid “cloud” in noisy environments. **A)** Trajectories of the hybrid states in the presence of random single-node perturbations given every 10 time steps for a (i) two-level and (ii) four-level hybrid state. The plots represent the position of the trajectories every 100 time steps. **B)** Density plots, showing the relative mean residence time (MRT) calculated over a simulation of 100 trajectories each for the top five hybrid steady states in (i) two-level and (ii) four-level formalism. Brighter (yellow) colors indicate higher MRT, while darker (blue) colors indicate lower MRT. **C)** Violin plots depicting the distribution of the phenotypic score across 5 X 100 trajectories for different intervals of noise application (x-axis), with (i) 5% and (ii) 50% perturbation at each interval

In contrast, four-level hybrid states largely maintained their hybrid phenotype, even after long-term stochastic simulations (**Figure 6A (ii)**, phenotypic score remains within the range *−* 1 and 1). Their trajectories were found primarily in regions corresponding to hybrid phenotypes with partial expression of both epithelial and mesenchymal nodes (**Figure 6B (ii)**). We further quantified this behavior across different perturbation intervals and magnitudes (i.e., number of nodes perturbed, **Figure 6C**). At high noise levels, that is short perturbation intervals (1 or 5) and with large perturbation magnitudes, two-level hybrid states also retained hybridness, primarily because the noisy environment prevented any phenotype from stabilizing. However, the distinction between two- and four-level states became clear at moderate and low noise levels: while two-level states localized readily to extreme phenotypic scores, four-level hybrid states consistently maintained intermediate scores (around 1 and -1). Even at long intervals between perturbations (100 timesteps), a fraction of four-level hybrids retained their hybridness.

We interpret this behavior as follows: four-level hybrid states, individually, are not more resistant to perturbations than two-level hybrids, as shown by their higher Hamming distances during transitions. However, they display collective stability as a “hybrid cloud,” which demonstrates significant resilience to noise.

Overall, our perturbation experiments highlight a key feature of four-level hybrid states: enhanced plasticity not observed in two-level hybrids. Although the stability of individual states under perturbation is comparable to that of two-level hybrids, the four-level model captures the dynamics of a hybrid cloud, a property with important implications for explaining the metastatic fitness of hybrid phenotypes.

## 3 Discussion

A central challenge in modeling complex biological systems such as cancer metastasis or drug resistance is that cell states which are relatively rare from a dynamical systems perspective can nonetheless play a dominant role in the biological outcome. In metastasis, for example, the vast majority of tumor cells that leave the primary site die either en route or upon arrival at the metastatic niche. The eventual metastatic lesion is founded by the exponential expansion of only a tiny, specialized subset of cells [39, 40]. This creates a fundamental tension for modeling approaches: whereas in many applications of Boolean network-based models the goal is to capture the typical behavior of the system [41, 42], in this context the rare states are the biologically relevant ones. As such, frameworks designed to describe average system behavior are insufficient on their own. Instead, metastasis and related processes demand models that are capable of capturing and predicting the emergence and stability of rare but functionally critical states.

Over the last decade, substantial experimental evidence has accumulated indicating that the key metastatic subpopulations exhibit a partial epithelial–mesenchymal transition (EMT) phenotype with substantially heterogeneous expression patterns, characterized by intermediate levels of both epithelial and mesenchymal gene expression [6, 9, 43]. This raises the critical question of whether Boolean models—which have been widely employed to study cell-fate decision networks [42]—can adequately represent such intermediate phenotypes. While some studies [21] demonstrated that Boolean networks could indeed generate a large number of hybrid E/M states, they corresponded to patterns in which subsets of EMT regulators were either fully “on” or “off”, rather than reflecting the graded expression levels observed in experimental data. Moreover, these spurious intermediate states were fragile, highly sensitive to noise, readily transitioning to epithelial or mesenchymal phenotypes [15]. These “metastable” hybrid phenotypes captured by previous Boolean models, therefore, cannot reliably carry on metastasis. Consequently, the partial EMT states that are most relevant to metastasis are largely inaccessible within conventional Boolean frameworks, due to the inherent limitation of reducing gene expression to binary on/off states. Our approach of multilevel Boolean formalism that started as a way capture the partial expression of E and M genes in hybrid states, goes on to highlight features of plasticity that make them metastatically relevant.

The theory underlying multi-valued networks has been developing in the recent past. Earlier implementations propose a multi-valued approach based on the number of downstream regulators a node has [44]. A recent implementation of this approach is the modeling formalism known as DSGRN [45, 46], a comprehensive treatment of the switching system that allows multiple expression levels for a given node based on the out-degree of the node. Using a generic system of ODEs, they define the range of expression levels each node can have and divide the range into multiple levels such that each level indicates active regulation of as many downstream genes. Such approaches have found success in describing the phenotypic landscape of small GRNs, but scale combinatorially with the network size. Another approach involves adding intermediate, fractional levels of expression such that the highest expression level remains 1. Several studies have aimed to understand the implications of expanding the state space in this manner [47–50], with the discovery of some interesting properties such as the multilevel formalisms seeing lesser instances of limit cycles, and capturing steady states using synchronous update scheme which in classic Boolean formalism could only be captured by asynchronous update schemes. From a biological standpoint, complexity has often followed context-specific functionality. A classic example is seen in signaling networks where proteins have different functions at different levels of phosphorylation [51]. Consequently, the selected nodes with the appropriate biological background allow multilevel expression.

Our current extension of Boolean formalism aims to explain the biological observation of hybrid phenotypes, while maintaining a formalism that does not require detailed biological information of such as the regulatory logic or kinetic parameters in the network. Using the multilevel Boolean model that allows two additional levels of expression (-0.5, 0.5), we capture hybrid steady states with a partial expression of E and M nodes. We find that these states have comparable SSF to the hybrid states obtained using the classic Boolean model, but have lower frustration due to their partial expression. A challenge for such a generic framework is to figure out the right level of granularity to be applied to the system. We found that increasing the number of levels beyond four (-1,-0.5, 0.5, 1) leads to loss of crucial properties of the biological phenotypes, namely, the dominant stability expected from terminal (epithelial and mesenchymal) phenotypes. Thus, we determined a four-level model to be appropriate for the current context. Using another theoretical modification of the Boolean formalism, we find that the hybrid states are predominantly generated due to the flexible expression of unregulated nodes during the simulation. When the effect of such nodes is nullified in the two-level Boolean model, hybrid states disappear completely. However, the multilevel model is capable of retaining the hybrid states, thereby indicating a topology driven mechanism of hybrid state emergence in the multilevel formalism.

The multilevel model enables three distinct features in the state-space as compared to the two-level model. 1) The number of unique stable hybrid states available increases significantly, with a majority of them having negligibly small basins of attraction. 2) The multilevel model allows for a reduced frustration of hybrid states, which can make them robust to noise, and 3) The multilevel hybrid states have enhanced stability as a collective, adding a new dimension of plasticity to hybrid states that is conducive to metastasis. The hybrid cloud-driven plasticity can allow metastatic cells to continue migrating collectively, while also changing their expression patterns that might allow for them to adapt to various stress factors. While the exact design principles of hybrid state-driven stemness and drug resistance are unclear, maintenance of hybrid state during metastasis, as allowed by the multilevel formalism, can help maintain these pro-survival phenotypes as well.

Our development of the multilevel formalism rooted in explaining hybrid E/M phenotypes. However, it can have broader applications. Team-based structure is found in GRNs of multiple cell-fate decision contexts. And in some of these contexts, hybrid states play a significant role. In the development, the hybrid characteristics (co-expression of markers corresponding to multiple cell fates) are often associated with progenitor cell types. High entropy has been utilized as the signature of the differentiability of a sample in transcriptomic data [52]. Hybrid states, especially the ones captured by the multilevel model, have high entropy. Similarly, in T cell differentiation, hybrid cells expressing markers of more than one class of T cells have been suggested to be precursors of different T cell types [53]. Thus, the multilevel Boolean formalism presented here can meaningfully be applied to a broader set of cell-fate decision networks.

## 4 Methods

### 4.1 Threshold-based models

In this manuscript, we used two kinds of models to simulate the EMP networks. The first is a class of multilevel threshold-based formalisms described by the equation below:

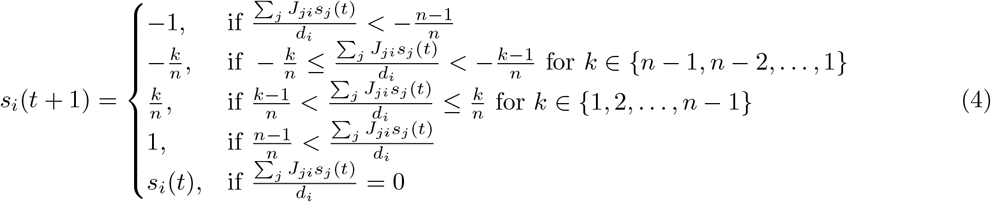

where *n* is the number of levels defining the state space granularity. We refer to these models as 2*n*-level models, with the two-level model being the classic Boolean formulation. *s*_*i*_(*t*) is the state of the *i*-th node at time *t, J*_*ji*_ is the regulatory interaction strength from node *j* to node *i*, and *d*_*i*_ is the in-degree of the *i*-th node. Note that the regulatory input *E* = ∑ _*j*_ *J*_*ji*_*s*_*j*_(*t*) *∈* [*−d*_*i*_, *d*_*i*_]. Since the expression levels are fractional, division by *d*_*i*_ ensures that the normalized input 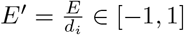.

The turn-off formalism is obtained by modifying the above equation as follows:

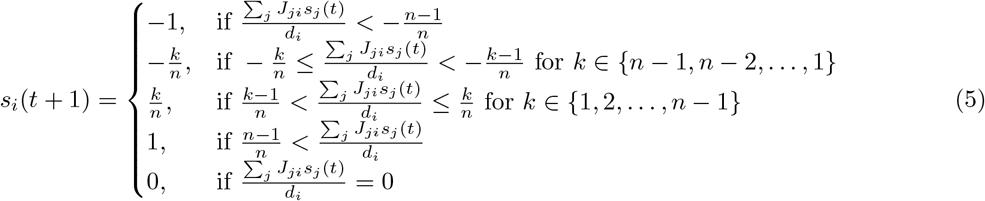

### 4.2 Network simulation

To simulate EMP network topologies, we first generated an ensemble of 10^7^ random initial conditions 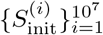, where each *S*^(*i*)^(0) is an *N* -dimensional vector with components randomly chosen from the discrete set 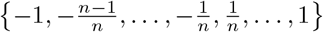, and *N* is the number of nodes in the network.

Using the appropriate update rules (Equations 4 and 5), each initial condition 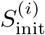 is updated asynchronously by randomly choosing one node per time step and updating the corresponding state component. The simulations continue until either a steady state is reached (i.e., *s*_*j*_(*t* + 1) = *s*_*j*_(*t*) for all *j ∈ {* 1, 2, …, *N }*) or a maximum of 1,000 time steps is reached. For the largest network (55N 111E), the time step limit is set to 10,000.

At the end of the simulations, we obtain an ensemble of steady states 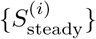. We then convert this ensemble into a frequency distribution by counting the fraction of initial conditions (out of 10^7^) that converge to each unique steady state.

### 4.3 Phenotypic score

For simulations, the phenotypic score of a steady state *S* is defined as:

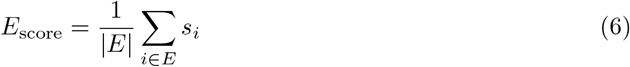

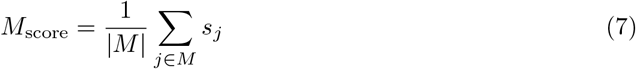

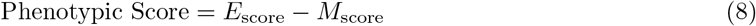

where *E* and *M* represent the sets of epithelial and mesenchymal nodes, respectively, and |*E*| and |*M* | are the cardinalities of these sets, i.e., the number of epithelial and mesenchymal nodes respectively.

### 4.4 Frustration

For a given state *S*, we define the frustration of an edge (*j, i*) as:

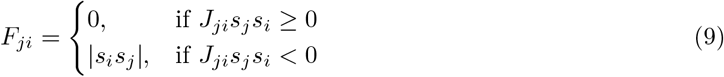

The total frustration of state *S* is defined as the normalized sum of frustrations over all edges, and is calculated as follows:

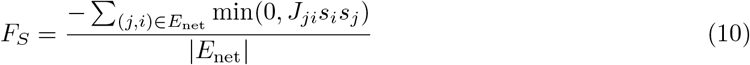

where *E*_net_ is the set of all edges in the network and |*E*_net_| is the total number of edges.

### 4.5 Transcriptomic data analysis

We obtained bulk RNA-seq data of carcinoma cell lines from the Cancer Cell Line Encyclopedia (CCLE) [32]. We extracted the expression matrix corresponding to an EMP gene set [25] from the cell line data. The expression matrix was log-normalized, followed by *z*-score normalization.

We calculated phenotypic scores using Equations 6–8, substituting the *z*-scores for *s*_*i*_, and further normalized the epithelial and mesenchymal scores to the range [*−* 1, 1]. For each gene, we defined “intermediate expression” as expression levels ranging from the 25th to 75th percentile of the gene’s expression vector across all samples. For each sample, we calculated the fraction of epithelial and mesenchymal genes exhibiting intermediate expression levels.

### 4.6 Perturbation analysis

We performed transient perturbations on the steady states of EMP networks and allowed the perturbed states to evolve for a fixed period. Each perturbation trajectory for a steady state *S* and perturbation level *k ∈ {*1, 2, …, *N }* was generated as follows:

1. Randomly choose a subset *K ⊂ {*1, 2, …, *N }* with |*K*| = *k*
2. Create a new initial perturbed state *S*^*′*^(0) where:

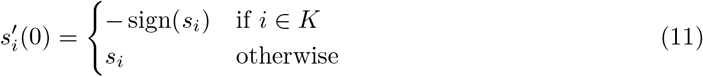
3. Simulate *S*^*′*^(0) using the appropriate update rules asynchronously, recording the state *S*^*′*^(*t*) every 5 time steps
4. Repeat steps 1–3 for multiple realizations until sufficient statistics are obtained

For single transient signal experiments (Figure 5), we simulated each perturbed state until convergence to a steady state. For noise experiments (Figure 6), we applied random single-node perturbations every 10 time steps and simulated for extended durations to observe long-term behavior.

## 5 Data and Code availability

The raw network files and the code used for network simulation, analysis and figure generation can be found at https://github.com/askhari139/MultilevelPaper. The simulations are performed using Julia and the corresponding code can be found at https://github.com/askhari139/Boolean.jl.

## 8 Author contributions statement

HL, ST and KH designed the research. KH, ST, and VA carried out the simulations. All authors discussed the results and participated in the preparation of the manuscript.

