## Supplementary Figures and Note for "A multilevel formalism to model the hybrid E/M phenotypes in Epithelial-Mesenchymal Plasticity"

### S1 Supplementary Methods

#### S1.1 Network randomization

To generate a random network, we take a permutation of the edge signs currently existing in the network. In other words, we took the third column of the edge table provided in Supplementary Table 1 and shuffled the entries of the column "Type". This ensures that the unsigned directed graph is preserved, thereby preserving all the connectivity-based graph properties such as the degree distributions, centrality measures, path connectivity and unsigned feedback loops. However, signed properties, particularly the abundance and connectivity of positive and negative feedback loops changes, which in turn effect the dynamics. The particular permutation in Figure S3 was chosen to ensure a mismatch of atleast half of the edge signs from the original network.

#### S1.2 Calculation of the fraction of weighted negative feedback loops

For each network, we identified all directed loops using the *simple_cycles* function of the *Networkx* module in *Python 3.10*. We then assigned a weighted score to each loop as the inverse of the number of edges participating in the loop. This weight penalizes longer loops, as they have a lesser chance of affecting the network dynamics. We then calculated the total score of positive loops (loops with an even number of inhibitory edges) and that of negative loops and calculated FWNC as the total score of negative loops divided by the total score of all loops.

#### S1.3 Threshold calculation for multi-level formalism

For multilevel models with large number of levels, the classification defined for the two- and four-level formalisms fails to capture terminal states. Hence, we defined a relative classification method by calculating a unique symmetric threshold for each state space. We scan potential threshold values from 0 to 2, with an interval of 0.05. For each threshold value, we define the phenotype to be terminal if $|Phenotypic\ score| > Threshold$ and hybrid otherwise. We then calculated the total frequency of the terminal and hybrid steady states. We picked the smallest threshold value that allows for the total frequency of terminal states to be higher than hybrid states.

#### S1.4 Calculation of frustration for CCLE data

To compare the simulation results with CCLE data, we calculated the frustration of the CCLE samples for the genes included in the 23N 89E EMP network. We eliminated the micro-RNAs except miR205, because their expression was not recorder in the CCLE samples. We then took the logarithm of the expression levels, and classified the samples as epithelial, mesenchymal or hybrid using the expressions and phenotypic scores similar to **Figure 1**. We then calculated the mean of the log expression for each gene across all samples and characterized the expression levels of the genes as $+1$ or $-1$ based on whether the value of the expression level is higher or lower than the corresponding mean. Using equation 10, we calculated the frustration of each CCLE sample. For comparison, we calculated a relative frustration for the CCLE data as well as the simulated data as the ratio of the frustration of each sample/steady state to that of the mean frustration of all hybrid samples/steady states.

### S2 Supplementary Note 1 : Inefficiency of transition into a continuous domain by further expanding the state space

Since increasing the granularity of the state space showed marked improvements in explaining biological data, we explored the effect of further expanding the state space. The corresponding update rules are given below:

$$S_{i_{(t+1)}} = \begin{cases} -1, & if \ \frac{\Sigma_j J_{ji} s_j}{d_i} < -\frac{n-1}{n} \\ -\frac{k}{n}, & if \ -\frac{k}{n} \leq \frac{\Sigma_j J_{ji} s_j}{d_i} < -\frac{k-1}{n} \quad \forall k \in n-1, n-2, ..., 0 \\ \frac{k}{n}, & if \ \frac{k-1}{n} < \frac{\Sigma_j J_{ji} s_j}{d_i} \leq \frac{k}{n} \quad \forall k \in 1, 2, ..., n-1 \\ 1, & if \ \frac{n-1}{n} < \frac{\Sigma_j J_{ji} s_j}{d_i} \\ S_{i_t}, & if \ \frac{\Sigma_j J_{ji} s_j}{d_i} = 0 \end{cases} \tag{12}$$

While we observed a significant increase in the number of steady states between 2-level and 4-level formalisms, the number seems to saturate as the number of levels increased, saturating at the order of $10^4$ for 23N 89E network (**Figure S7A**). This held true for other networks as well, albiet less pronounced for 16N 32E network (**Figure S14A(a), B(a) and S15A(a)**). As the number of levels increases, we notice a consistent decrease in the range of phenotypic scores of the steady states. While the maximum score of two-level formalism was 2, the same for the twenty-level formalism was close to 1 (**Figure S7B, S14A(b), S7B(b), and S15B(b)**). Correspondingly, none of the states have a maximum expression of all epithelial or mesenchymal nodes. Therefore, the multilevel model does not capture any true "terminal" state. Furthermore, the stability of the states at the ends of the phenotypic score spectrum, representative of the maximum extent of epithelial and mesenchymal phenotype, significantly goes down even for 6-level models. However, states with high stability still appear, with more than 80% of the state space converging to the first 10 steady states ordered by frequency for 23N 89E and 15N 60E networks (**Figure S7C, S14A(c)**). For the 16N 32E and 27N 76E networks showed approximately 50% states converging to the first 10 steady states (**Figure S14B(c), S15A(c)**). Applying the previous classification with re-scaling the phenotypic scores also would not yield a reasonable classification, because the steady states near the boundaries of the phenotypic score have low frequency (**Figure S7B**).

To capture the effectively terminal states in the multi-level formalisms, we adopted a new system of steady-state classification. First, we clearly see a need to normalize the phenotypic score of these steady states as we increase the number of levels in the model. Hence, we divided the epithelial and mesenchymal scores by the maximum magnitude of the corresponding score observed across all steady states, such that the range of scores now becomes -1 to 1. Second, we decided to relax the constraints on the hybrid classification system. Note that the samples classified to be in epithelial or mesenchymal states have a wider range of scores. As two-level and four-level models have been largely discrete compared to the experimental data, each expression level allowed in these models covers a large range of expressions on a continuous scale, validating the strict definitions that we applied to classify the states. However, as the number of levels increase, we move closer to the continuous scale of expressions in our model, losing the discrete nature of the model. Thus, we decided to relax our classification criteria to identify terminal states by assigning a threshold to the E and M scores such that states with an E score $>$ threshold and an M score $< 1 - *$threshold are considered epithelial and vice versa. To calculate the threshold, we scanned through the range of possible thresholds from 0 to 1. For each value, we characterized the state space into terminal and hybrid states, and asked if the total frequency of terminal states is greater than the total frequency of hybrid states, a property consistently observed in EMP experiments. We used the minimum (**Figure S7D**). Furthermore, we devised an additional condition to identify this threshold for a given network-level combination: the total frequency of hybrid states must be less than the total frequency of terminal states, in accordance with experimental observations [6]. Although we did not restrict the SSF of individual steady states, our new classification method retained a higher average SSF for terminal states across all networks (**Figure S7E, S14, S15**).

As the number of levels increases, the multilevel model fails to capture the terminal and incomplete terminal states, wrongly classifying all steady states as hybrid. Furthermore, the steady states with extreme phenotypic score, which should be closest to epithelial and mesenchymal phenotypes, have lower steady state frequency (abundance) than the steady states in the middle of the phenotypic score spectrum. Clustering the expression levels of the steady states and then classifying the clusters into terminal and hybrid phenotypes based on the E and M scores (compare the mean E scores and M scores of the clusters and classify the clusters with the highest E score or highest M score as terminal and the remaining as hybrid – similar to the classification of samples in CCLE data in Figure 1) also did not capture the expected stability differences between terminal and hybrid states consistently. Considering the classification method as a part of the model, which can be tuned to fit expected behavior, we defined an adaptive threshold for classifying the steady states, such that any state with a phenotypic score greater than the threshold or less than -1*threshold is classified as terminal, and other states are classified as hybrid. The estimation of this threshold was dependent on the condition that the total frequency of terminal states must be higher than that of hybrid states. While this threshold fits the expected pattern by design, the resultant terminal and hybrid states are not well separated in the E score- M score space. Given these drawbacks of not being able to capture distinct terminal states with the expected stability properties, we decided to limit the expansion of state space granularity to four levels.

### S3 Supplementary Note 2 : Theoretical limits of frustrations for the stability of a state

An edge $ji$ is said to be frustrated for a given state $S(t)$ (not necessarily steady state), if

$$J_{ji}s_i(t)s_j(t) < 0 \tag{13}$$

Correspondingly, the frustration of a state is defined as follows:

$$\begin{aligned} f_S(t) &= \frac{\sum_i \sum_j \mathbb{1}_{\mathbb{Z}^-}(J_{ji}s_i(t)s_j(t))}{\sum_i \sum_j |J_{ji}|}, \quad further \\ \sum_i \sum_j |J_{ji}| &= \sum_i T_{1_i} + T_{2_i} \quad where \\ T_{1_i} &= \sum_j \mathbb{1}_{\mathbb{Z}^-}(J_{ji}s_i(t)s_j(t)) \\ T_{2_i} &= \sum_j \mathbb{1}_{\mathbb{Z}^+}(J_{ji}s_i(t)s_j(t)) \end{aligned} \tag{14}$$

Thus,

$$f_{S(t)} = \frac{\sum_i T_{1_i}}{\sum_i T_{1_i} + T_{2_i}} \tag{15}$$

In the ising boolean formalism, the update rules are defined as follows:

$$s_i(t+1) = \begin{cases} +1, \sum_j J_{ji}s_j(t) > 0 \\ -1, \sum_j J_{ji}s_j(t) < 0 \\ s_i(t), \sum_j J_{ji}s_j(t) = 0 \end{cases} \tag{16}$$

with standard definitions of all involved terms. The update rules indicate that, for the activity of a node to remain conserved in the next time step,

$$s_i(t+1) = \begin{cases} \sum_j J_{ji}s_j(t) >= 0, s_i(t) = +1 \\ \sum_j J_{ji}s_j(t) <= 0, s_i(t) = -1 \end{cases} \tag{17}$$

In other words,

$$s_i(t+1) = s_i(t) \ if \ \sum_j J_{ji}s_j(t)s_i(t) >= 0 \tag{18}$$

Thus, the condition for a state $S(t)$ to be a steady state is:

$$\sum_j J_{ji}s_j(t)s_i(t) >= 0 \forall i \tag{19}$$

It is possible to establish a relationship between frustration and the possibility of a state being stable. Now, consider the following two extreme cases of states:

- All edges are frustrated, i.e., $Frustration = 1$. In this case, Since $J_{ji}s_i(t)s_j(t) < 0$ for all $i, j$, the condition in 18 is never satisfied. Hence, none of the nodes in the state can retain their activity, and therefore the state cannot be a steady state.
- No edge is frustrated, i.e., $Frustration = 0$. In this case, the condition in 18 is satisfied for all $i, j$ and hence for all nodes, making the state steady.

These extreme cases hence seem to indicate that the higher the frustration, the lesser the chance of a state being steady. However, having the frustration value as 1 is seldom possible, due to the presence of signal nodes (not influenced by any other node in the network and therefore can retain any activity), negative feedback, and feed-forward loops. Thus, there exists a threshold value of frustration above which a state ceases to be a steady state.

Note that the magnitude of the term $J_{ji}s_i(t)s_j(t)$ is always 1. Therefore, the condition for the stability of the $S(t)$ (Equation 19) can be re-written as follows:

$$\begin{aligned}
T_{1_i} - T_{2_i} &\leq 0 \forall i \\
i.e.,\ T_{1_i} &\leq T_{2_i} \forall i \\
i.e.,\ \sum_i T_{1_i} &\leq \sum_i T_{2_i} \\
\frac{\sum_i T_{2_i}}{\sum_i T_{1_i}} &\geq 1
\end{aligned} \tag{20}$$

Rewriting 15, we get

$$f_{S(t)} = \frac{1}{1 + \frac{\sum_i T_{2_i}}{\sum_i T_{1_i}}} \tag{21}$$

Note that $\sum_i T_{1_i} = 0$ ensures that there are no negative elements in LHS of Equation 19 and thus ensures the stability of $S(t)$. Similarly, this condition sets $f_{S(t)} = 0$. Hence, in the above equations, we can use $\sum_i T_{1_i} \neq 0$ without loss of generality. Using 20, we have

$$\begin{aligned}
\frac{1}{f_{S(t)}} &= 1 + \frac{\sum_i T_{2_i}}{\sum_i T_{1_i}} \geq 2 \\
\therefore f_{S(t)} &\leq 1/2
\end{aligned} \tag{22}$$

Thus, Equation 22 defines the necessary condition for a state to be stable in the ising formalism.

### S4 Supplementary Figures

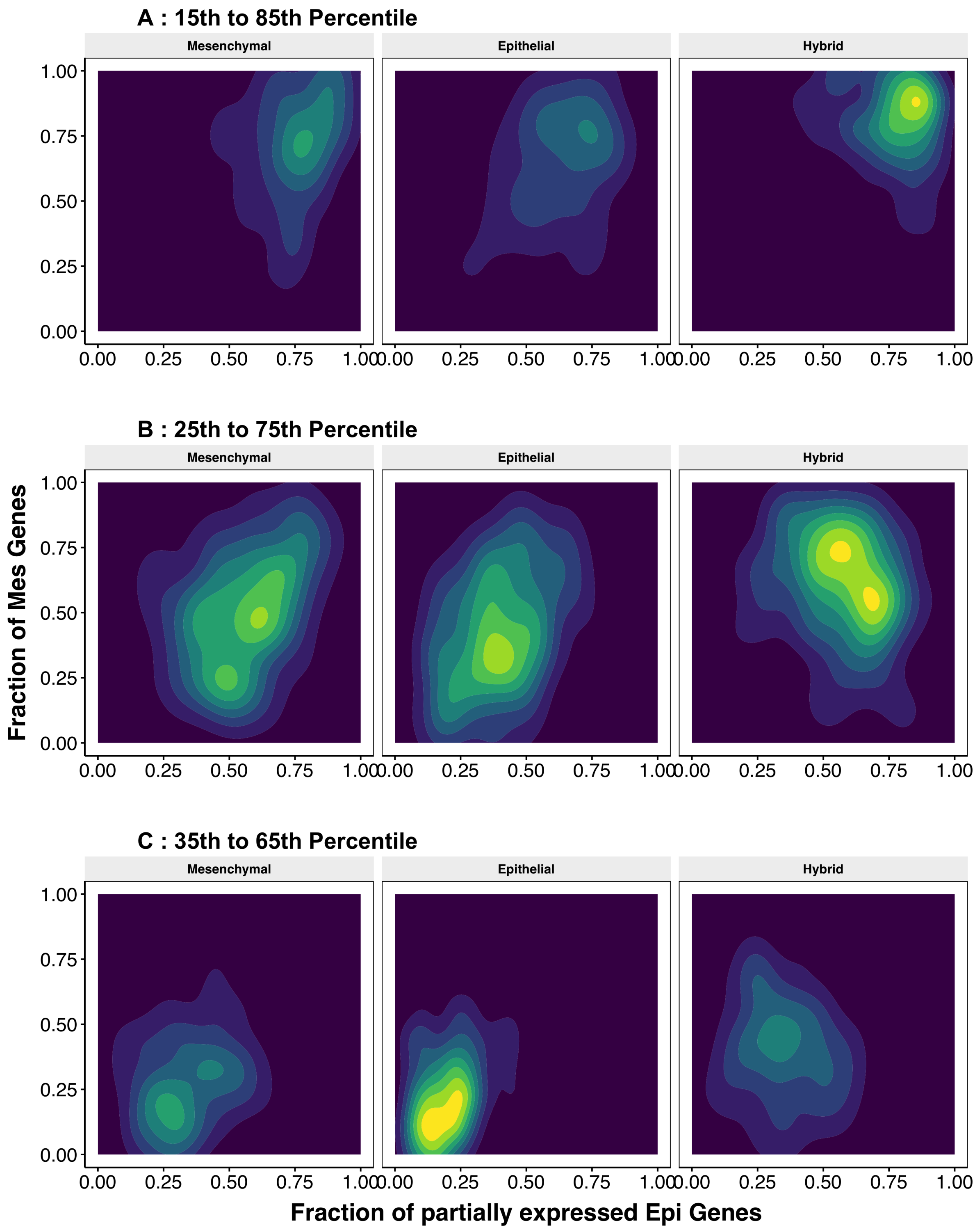


Figure S1: Testing different ranges to designate partial expression of a gene in a sample. The ranges are given along with the label for each row. Each row has three panels, corresponding to epithelial, mesenchymal and hybrid samples. Each panel is a density plot, with the yellow regions marking the region with the highest concentration of samples. The x and y axis map the fraction of epithelial and mesenchymal genes with partial expression, respectively.

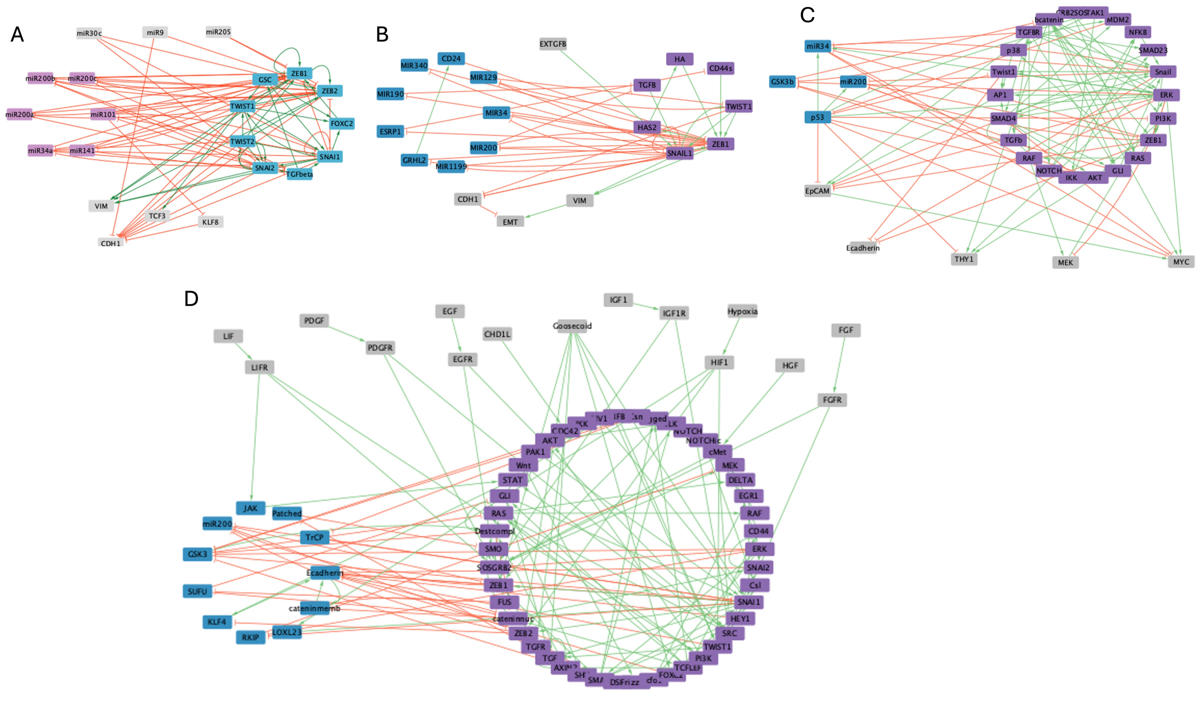

Figure S2: Wiring diagrams of the networks studied in this manuscript: **A)** 22N 82E, **B)** 20N 40E, **C)** 31N 95E and **D)** 70N 142E. In each diagram, the blue nodes are mesenchymal, pink are epithelial. Grey nodes at the top of any diagram are input nodes, while those at the bottom are output nodes. Note that in some cases, secondary input/output nodes exist that only have incoming/outgoing edges from another input/output and have also been considered as input/output. Examples: EpCAM in **D** and PDGFR in **E**. The network diagrams were generated using Cytoscape [54].

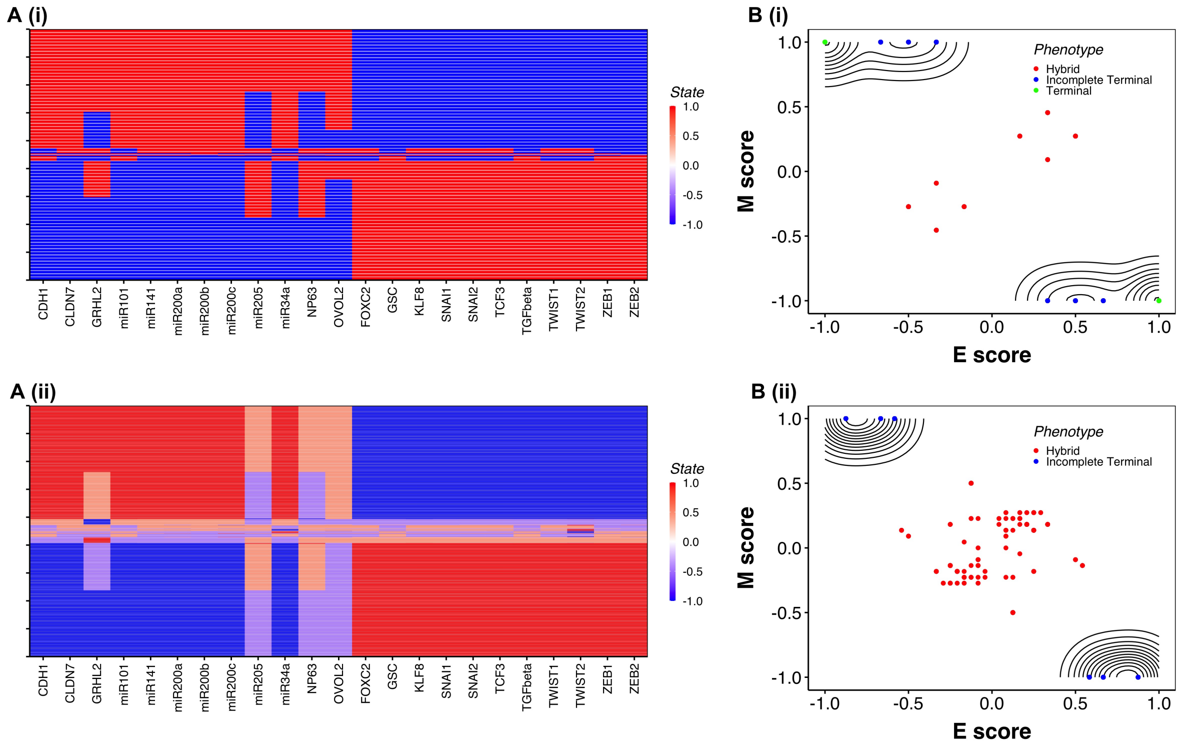


Figure S3: **A**Heatmaps and **B**contour plots depicting the ensemble of steady states obtained from the $10^7$ initial conditions for 23N 89E network with **(i)**two-level and **(ii)**four-level model.

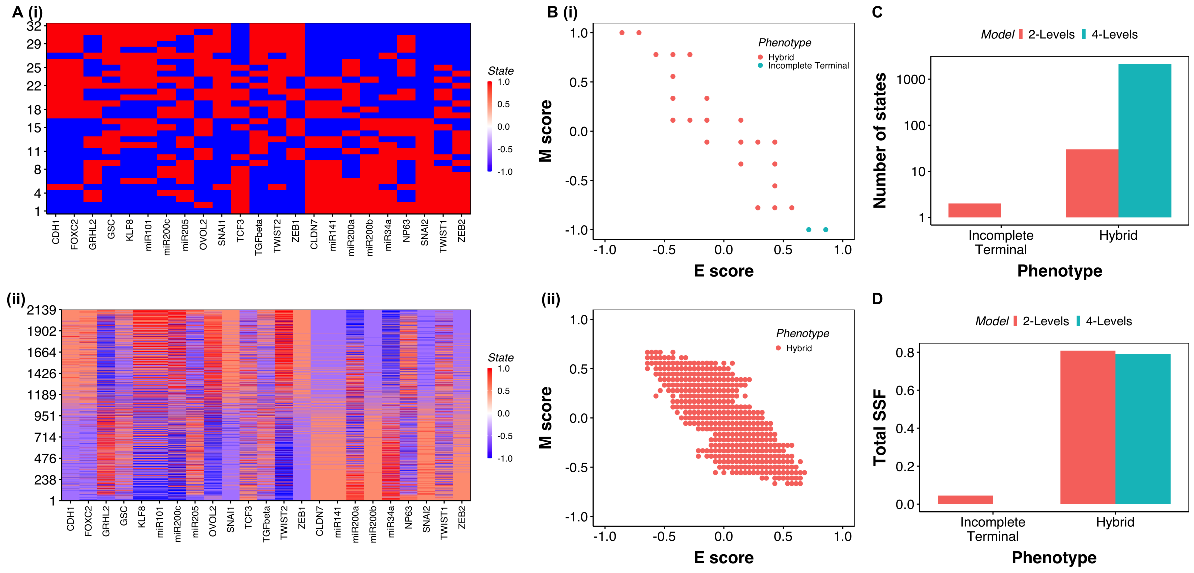


Figure S4: Composition and characteristics of steady states obtained from 2-level and 4-level model simulations of a network generated from the 23N 89E network by randomizing the edges. Panels are arranged in the same way as **Figure 2**

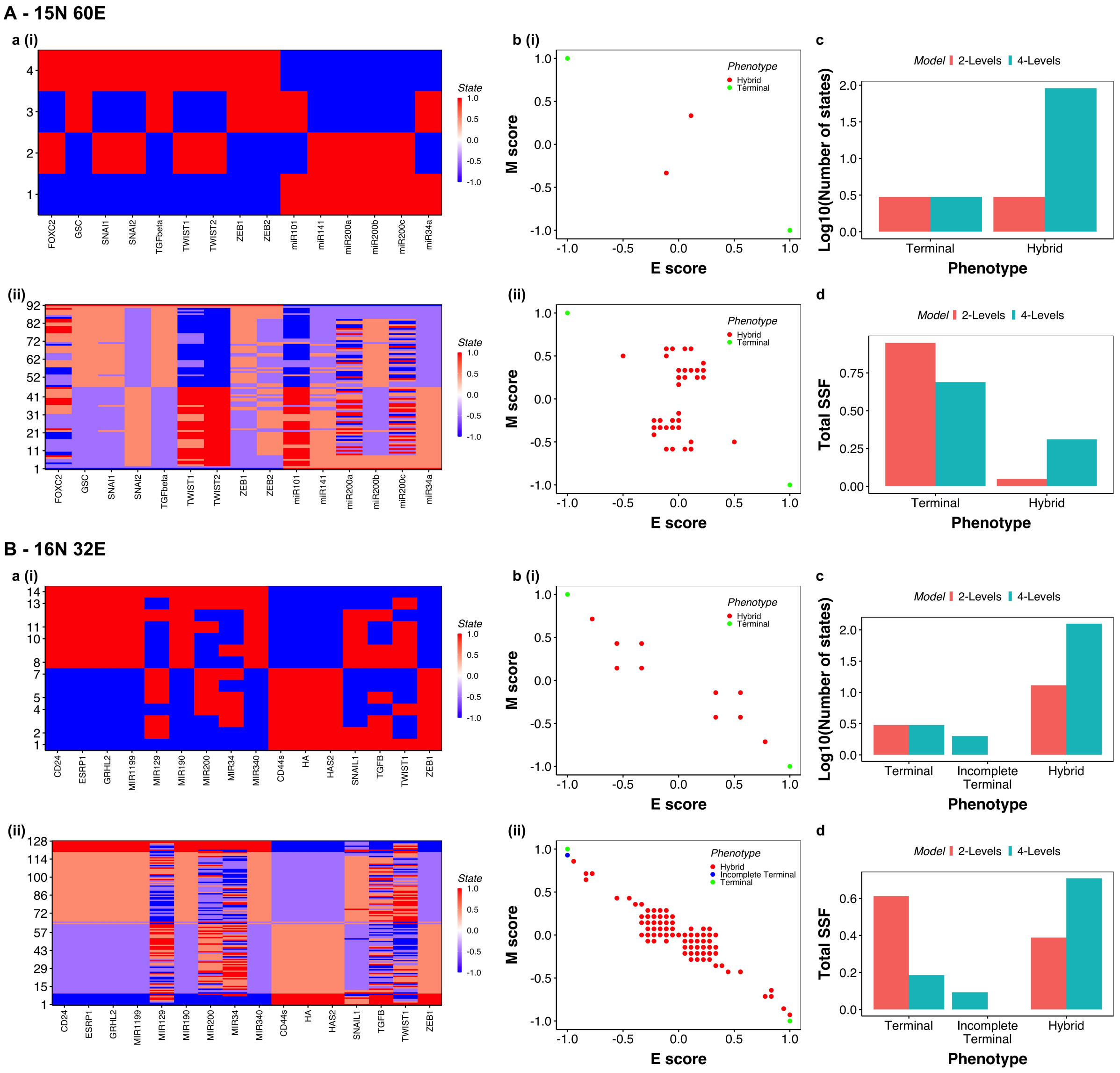


Figure S5: Composition and characteristics of steady states obtained from 2-level and 4-level model simulations of **A** 15N 60E[24] network and **B** 16N 32E[22] network. Each panel is arranged in the same way as **Figure 2**

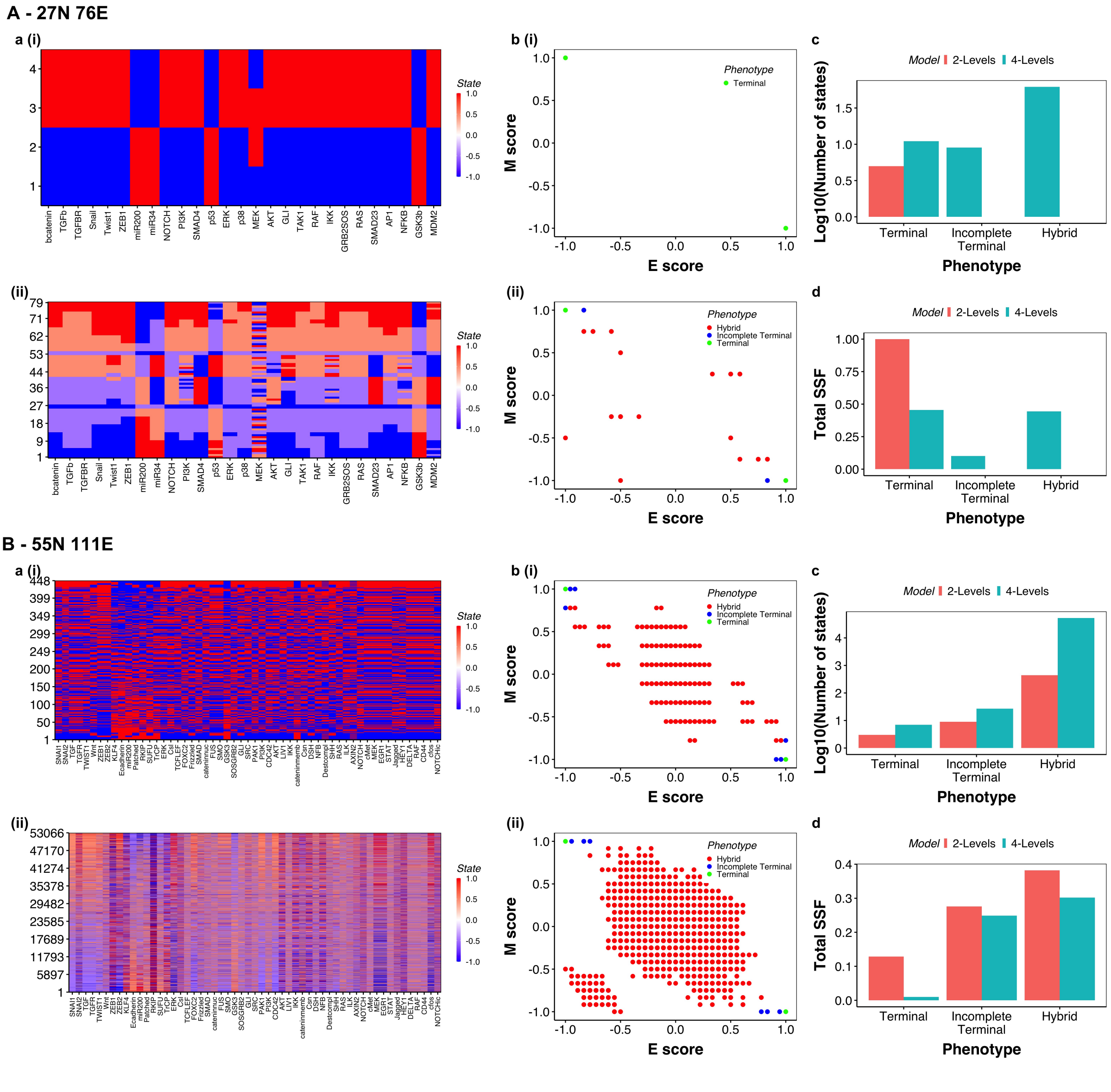


Figure S6: Composition and characteristics of steady states obtained from 2-level and 4-level model simulations of **A** 31N 95E[23] network and **B** 57N 113E[21] network. Each panel is arranged in the same way as **Figure 2**

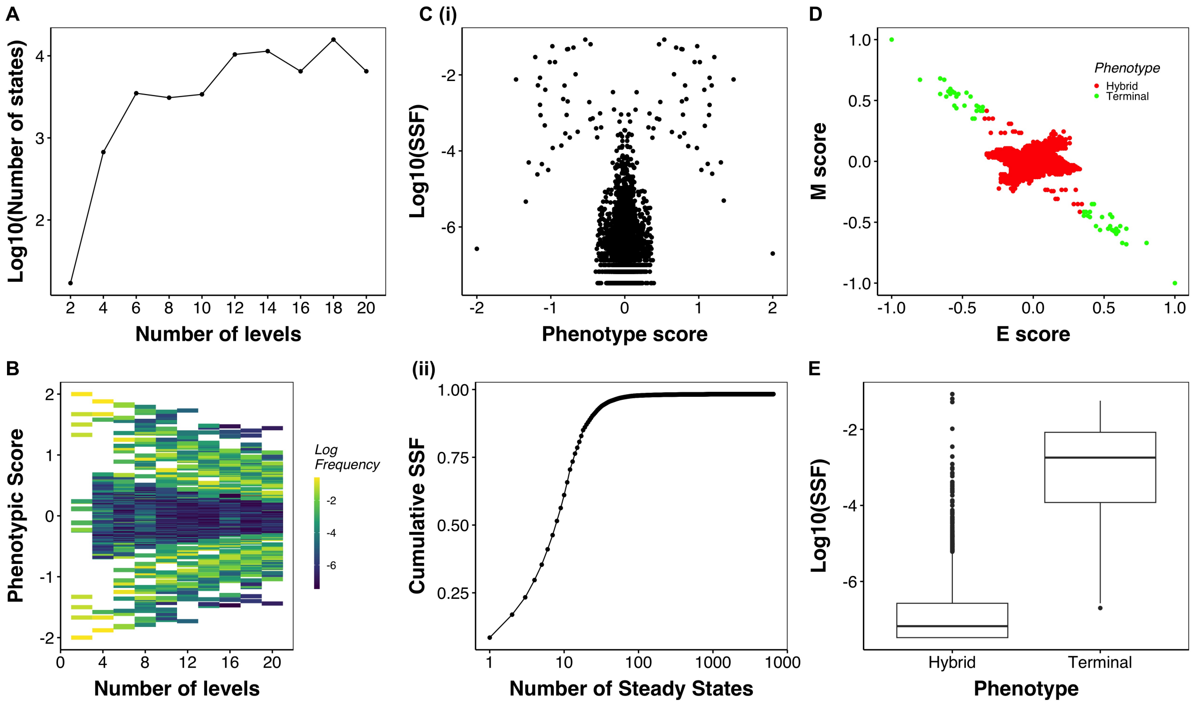


Figure S7: **Increased number of levels in the multi-level model narrows down the range of phenotypic score A** For multilevel formalisms of different numbers of levels, the number steady states. **B** Segment plot depicting the phenotypic score of the steady states with increasing number of levels in the multi-level model. **C** (i) 2-D contour plot of phenotypic score vs SSF for steady states of 23N90E network simulated using 20-level formalism. (ii) Scatterplot depicting the cumulative SSF (y-axis) of the n most frequent steady states (x-axis). **D** Scatterplot depicting the classification of steady states into terminal and hybrid states using the relative state classification system. **E** Steady state frequency of states clustered by the normalized phenotypic score.

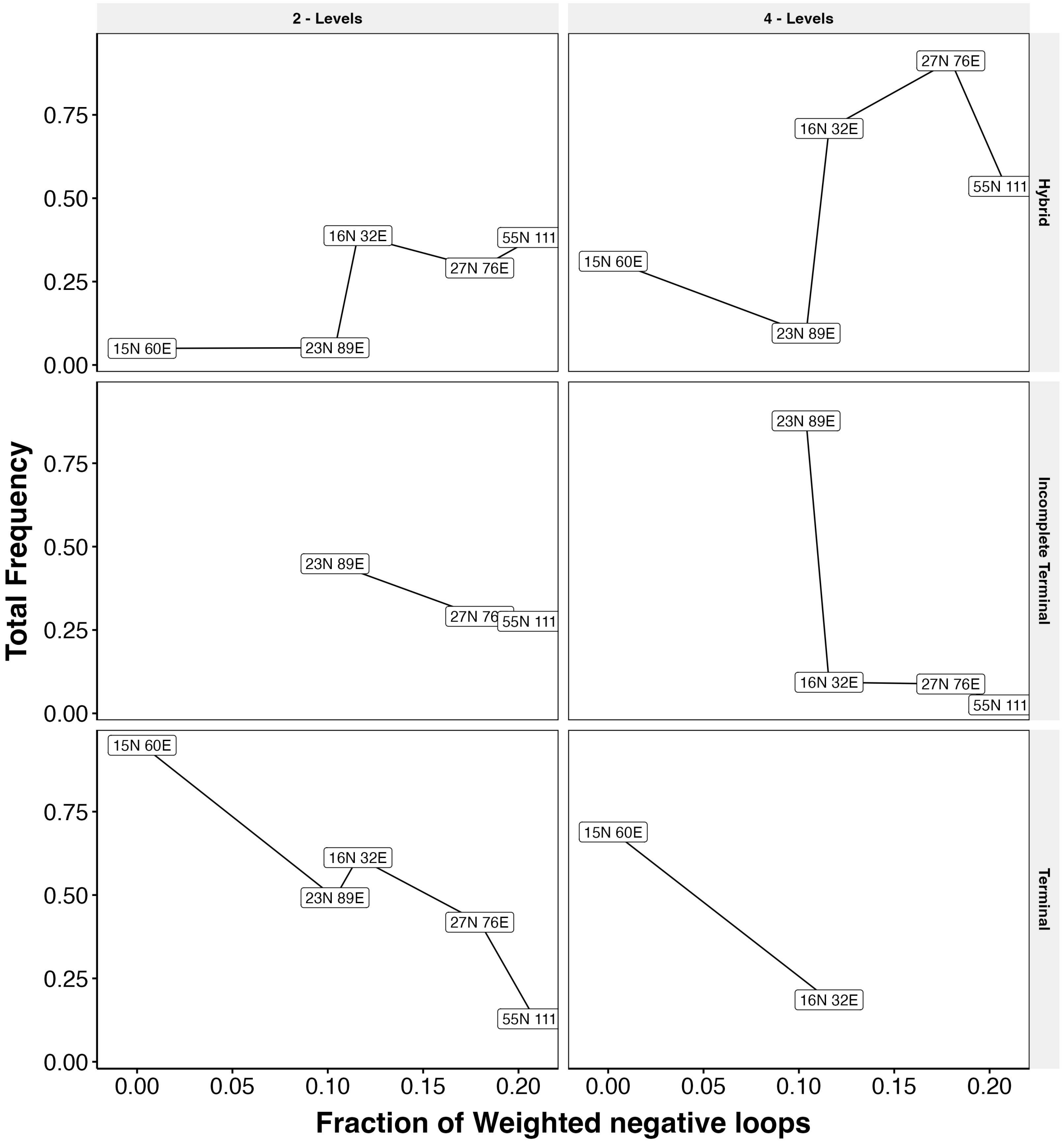


Figure S8: Line plots showing the change in frequency of hybrid, terminal, and incomplete terminal states with change in the fraction of weighted negative feedback loops. Network names are labeled with each point

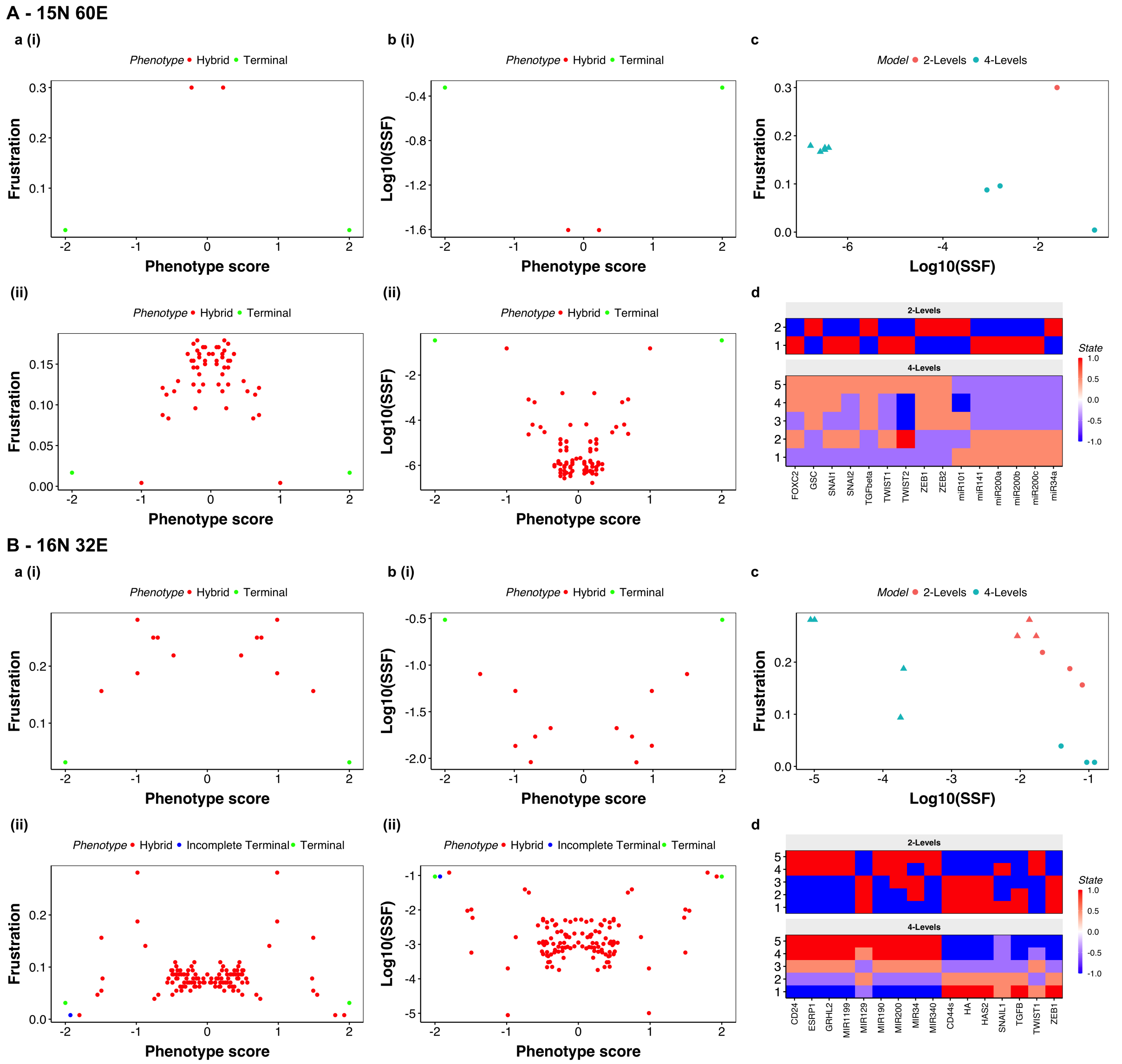


Figure S9: Composition and characteristics of steady states obtained from 2-level and 4-level model simulations of **A** 15N 60E[24] network and **B** 16N 32E[22] network. Each panel is arranged in the same way as **Figure 3**

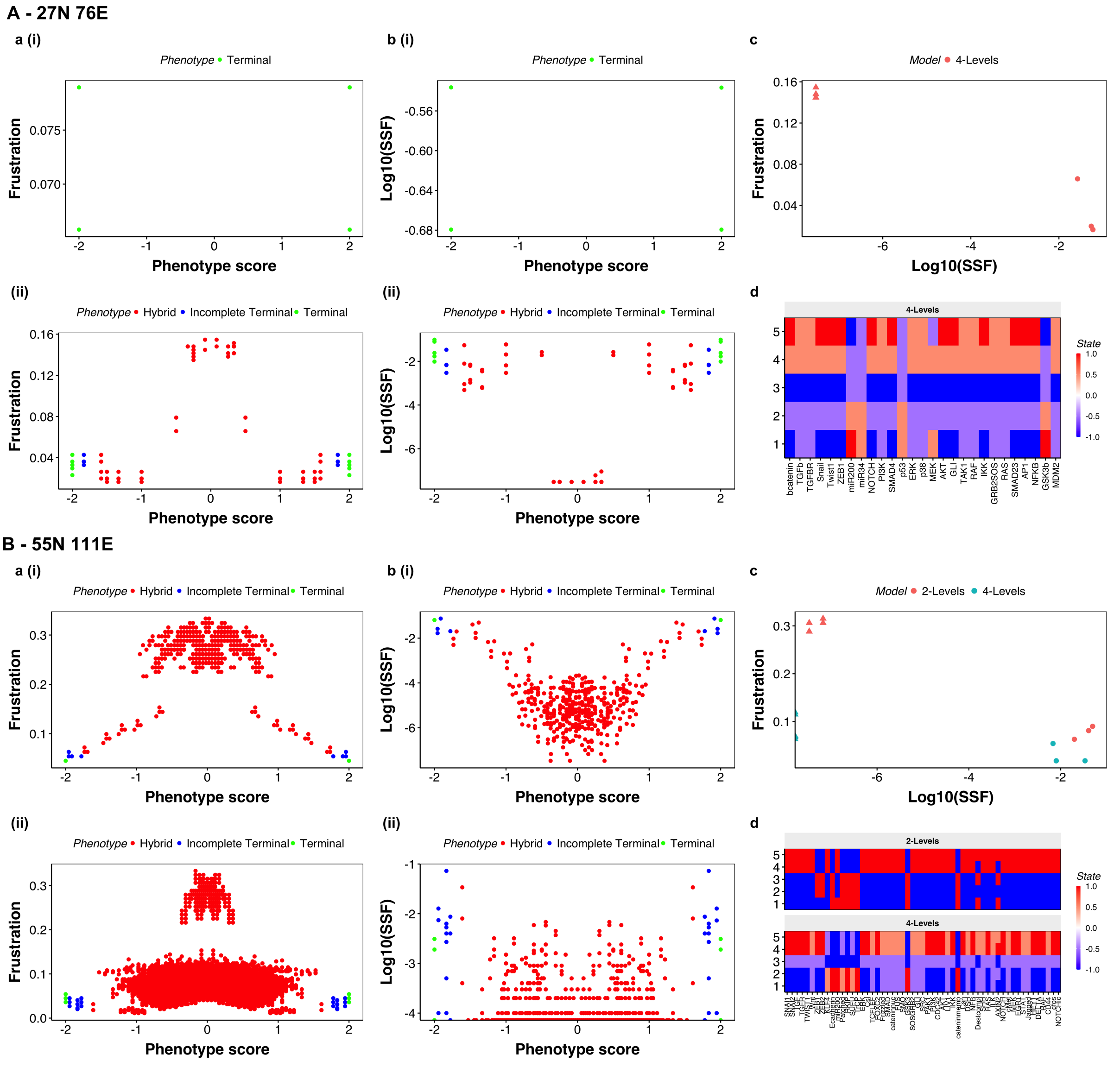


Figure S10: Composition and characteristics of steady states obtained from 2-level and 4-level model simulations of **A** 31N 95E[23] network and **B** 57N 113E[21] network. Each panel is arranged in the same way as **Figure 3**

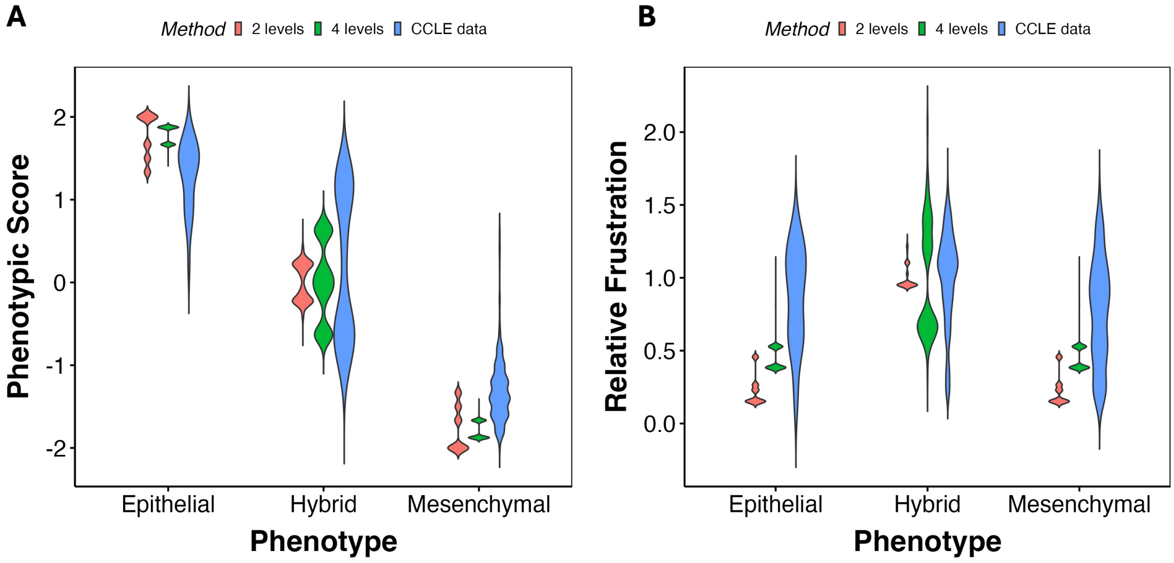


Figure S11: Comparison of simulation results with biological data. **A** Phenotypic scores of the expression levels of 23N 89E network nodes in CCLE data, in comparison with the two and four level model simulations. **B** Frustration calculated by discretizing the expression levels of EMT genes in CCLE around their mean and using the 23N 89E network topology with the discretized data.Relative frustration, obtained by dividing the frustration of each sample with the mean of hybrid samples, is compared to the relative frustration of two and four level simulations.

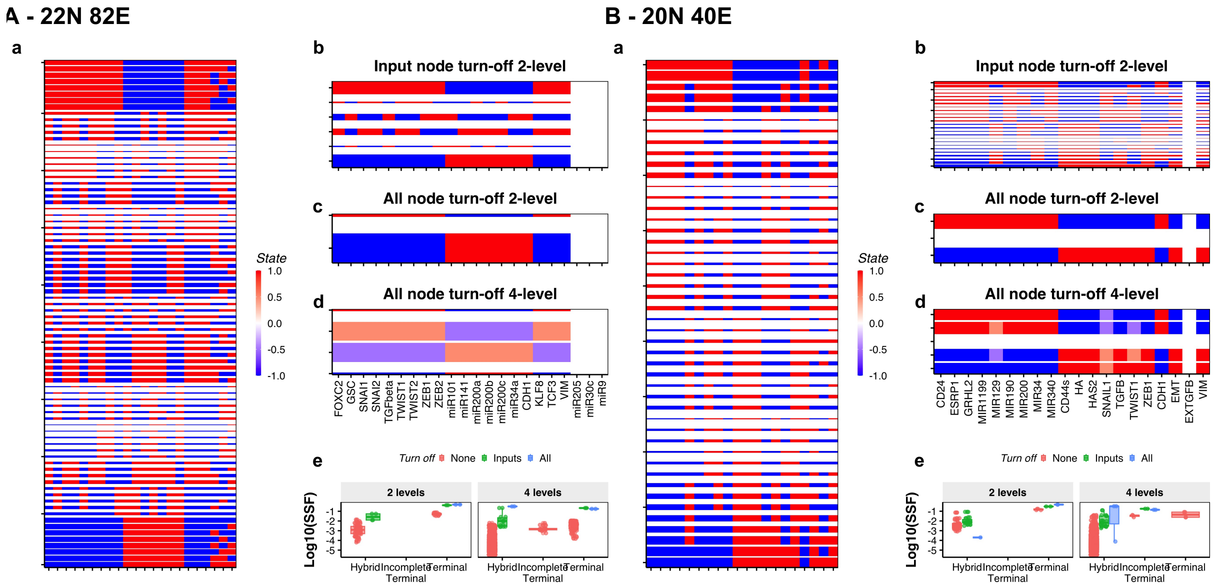


Figure S12: Turn off formalism applied to **A** 15N 60E[24] network and **B** 16N 32E[22] network. Each panel is arranged in the same way as **Figure 4**

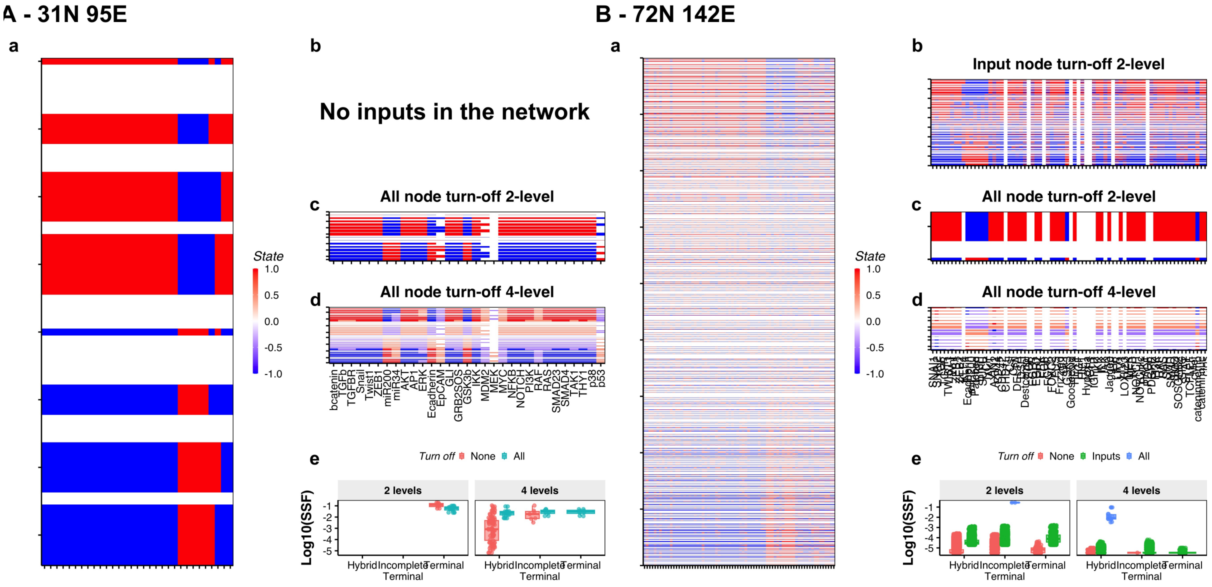


Figure S13: Turn off formalism applied to **A)** 31N 95E[23] network and **B)** 57N 113E[21] network. Each panel is arranged in the same way as **Figure 4**

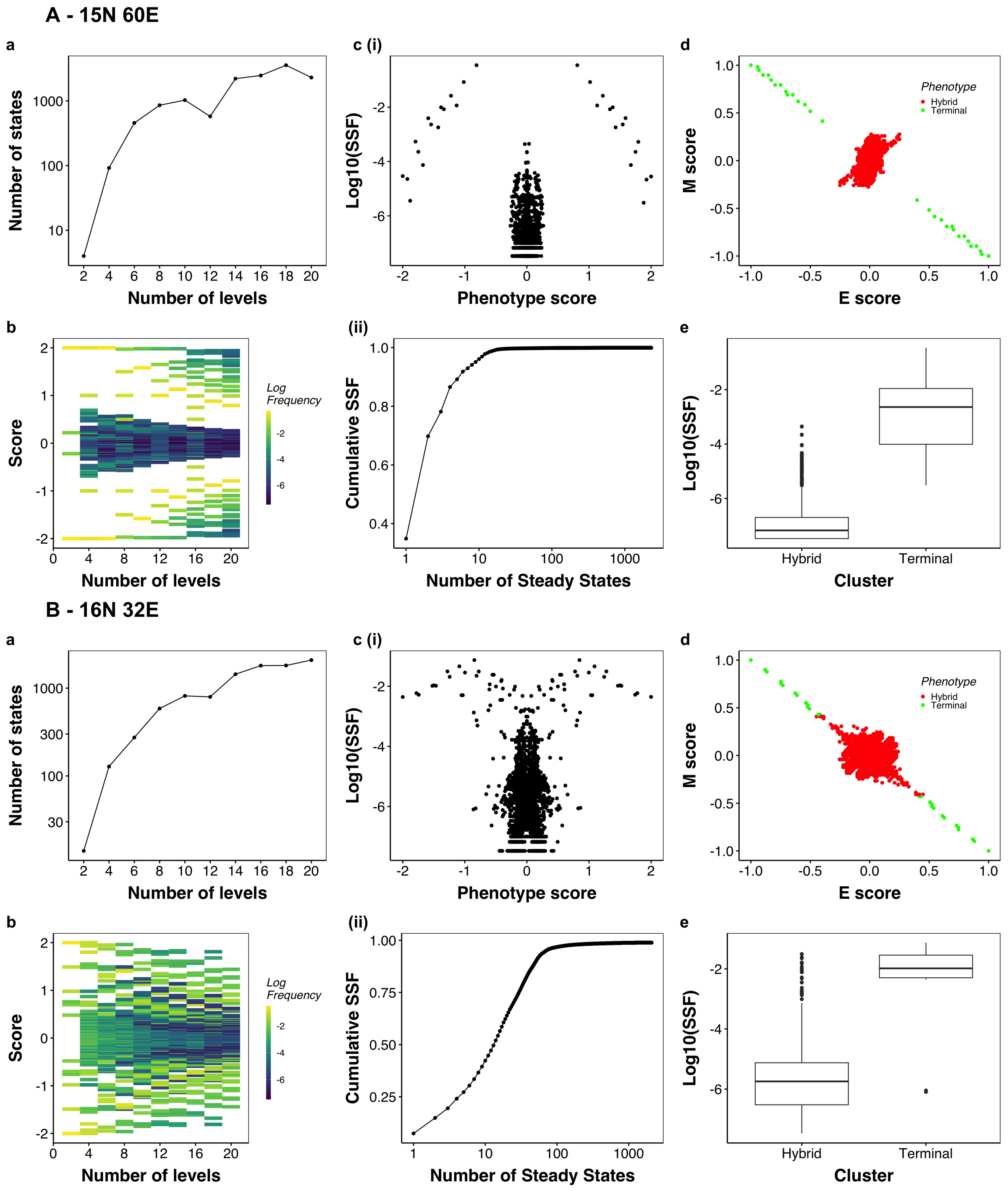


Figure S14: Analysis of multilevel model simulations for **A** 15N 60E[24] network and **B** 16N 32E[22] network. Each panel is arranged in the same way as **Figure S7**

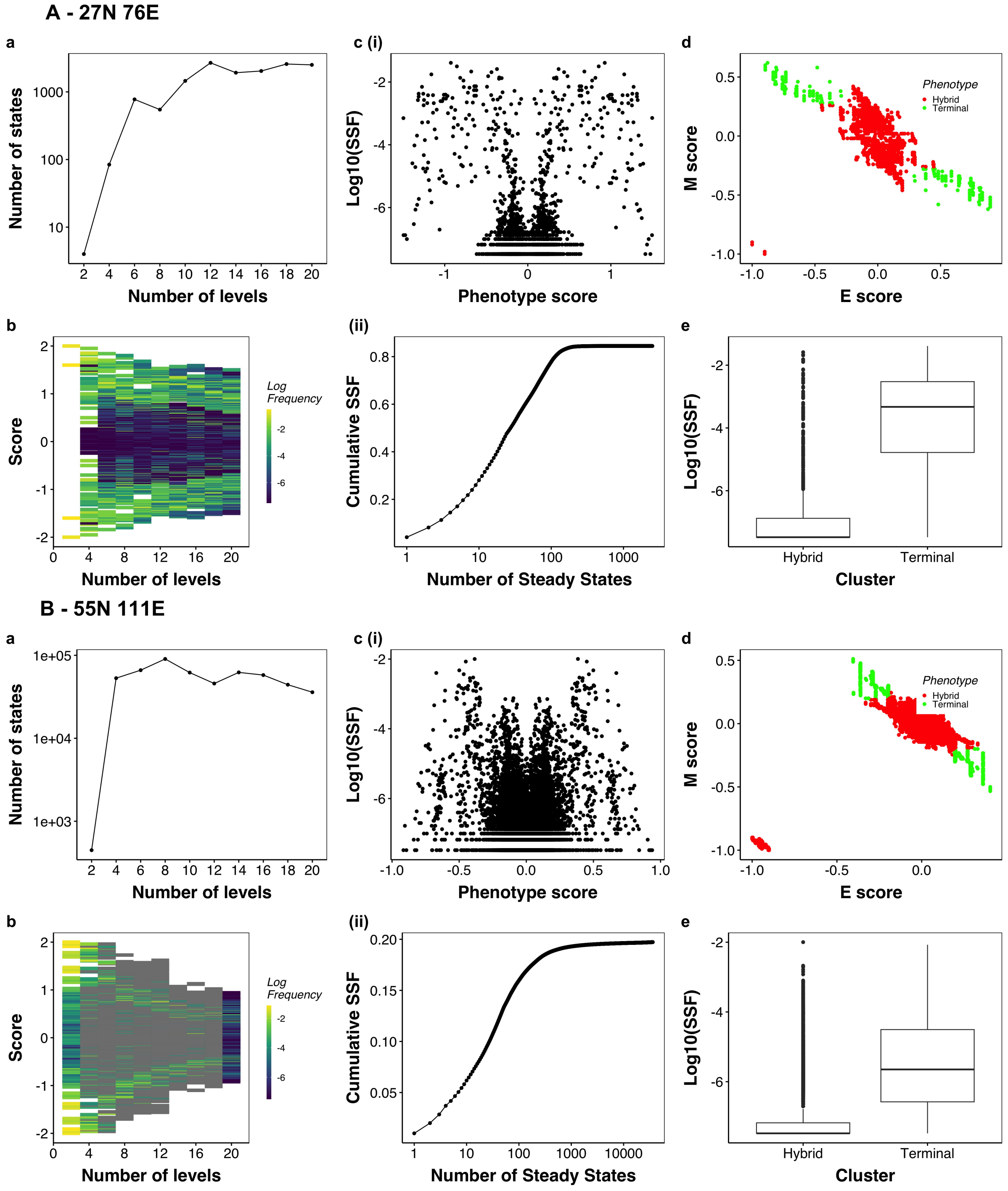


Figure S15: Analysis of multilevel model simulations for **A** 31N 95E[23] network. Each panel is arranged in the same way as **Figure S7**

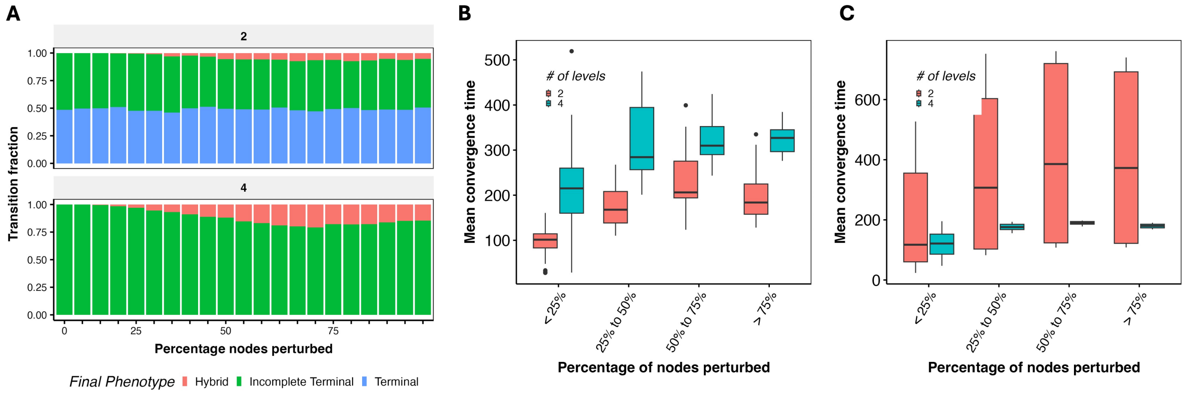


Figure S16: Transient perturbations of terminal steady states of 23N 89E network. **A** Transition probabilities of terminal states for two and four level models as a function of the extent of perturbation. **B** Time taken for convergence for transitions between hybrid states **C** Same as **B** but for transitions within terminal and incomplete terminal states.

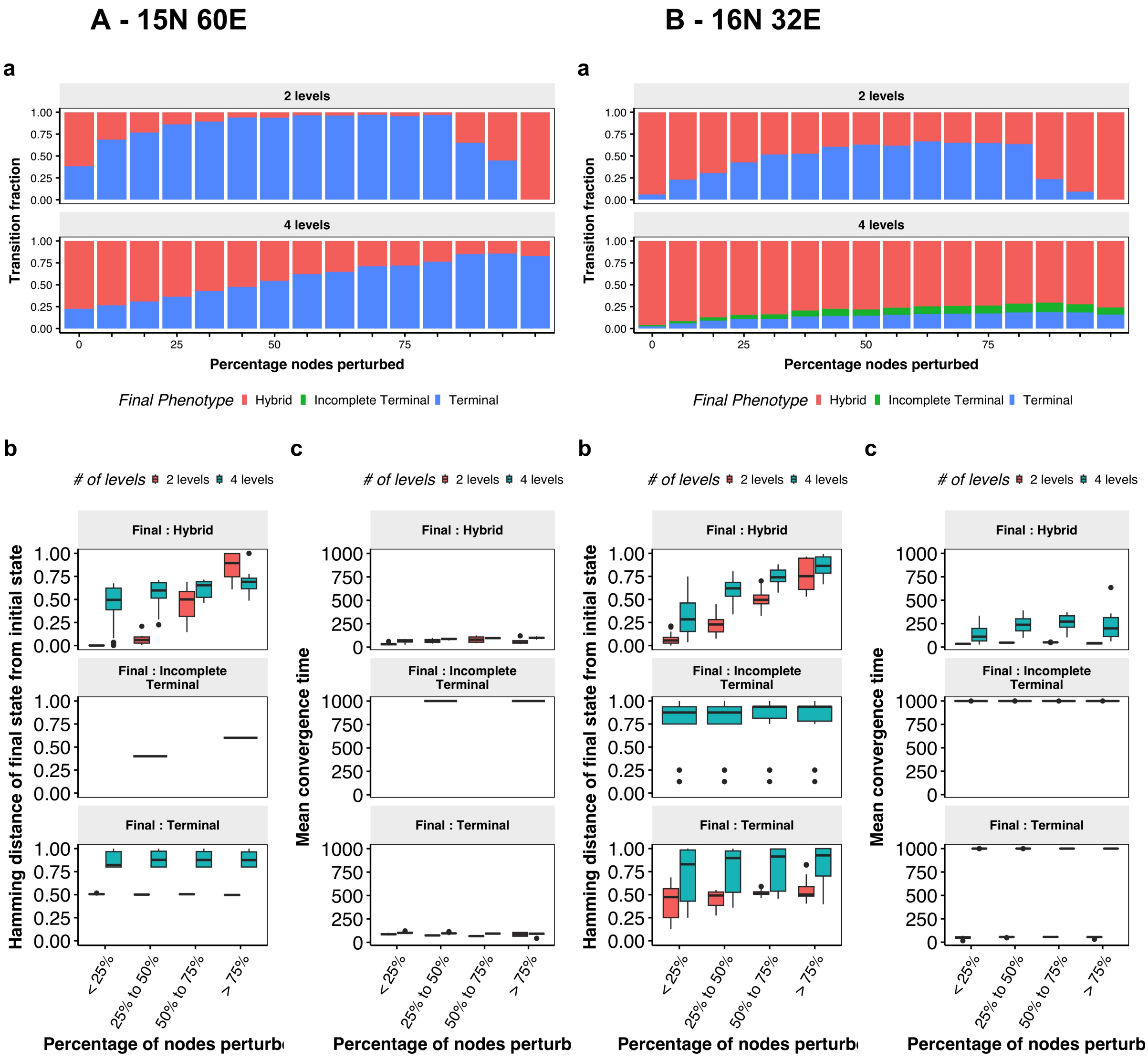


Figure S17: Analysis of multilevel model simulations for **A** 15N 60E[24] network and **B** 16N 32E[22] network. The subpanel details are as follows: **a** Transition probability from hybrid states to other phenotypes for two and four-level formalism. Note that the increase in transition to hybrid phenotype in two-level model after 75% perturbation is an artefact of the symmetry of threshold-based formalism. **b** Boxplots depicting the hamming distance between the initial and final steady state, with each panel showing classified by the phenotype of the final state. **c** Same as b, but shows the time to converge.

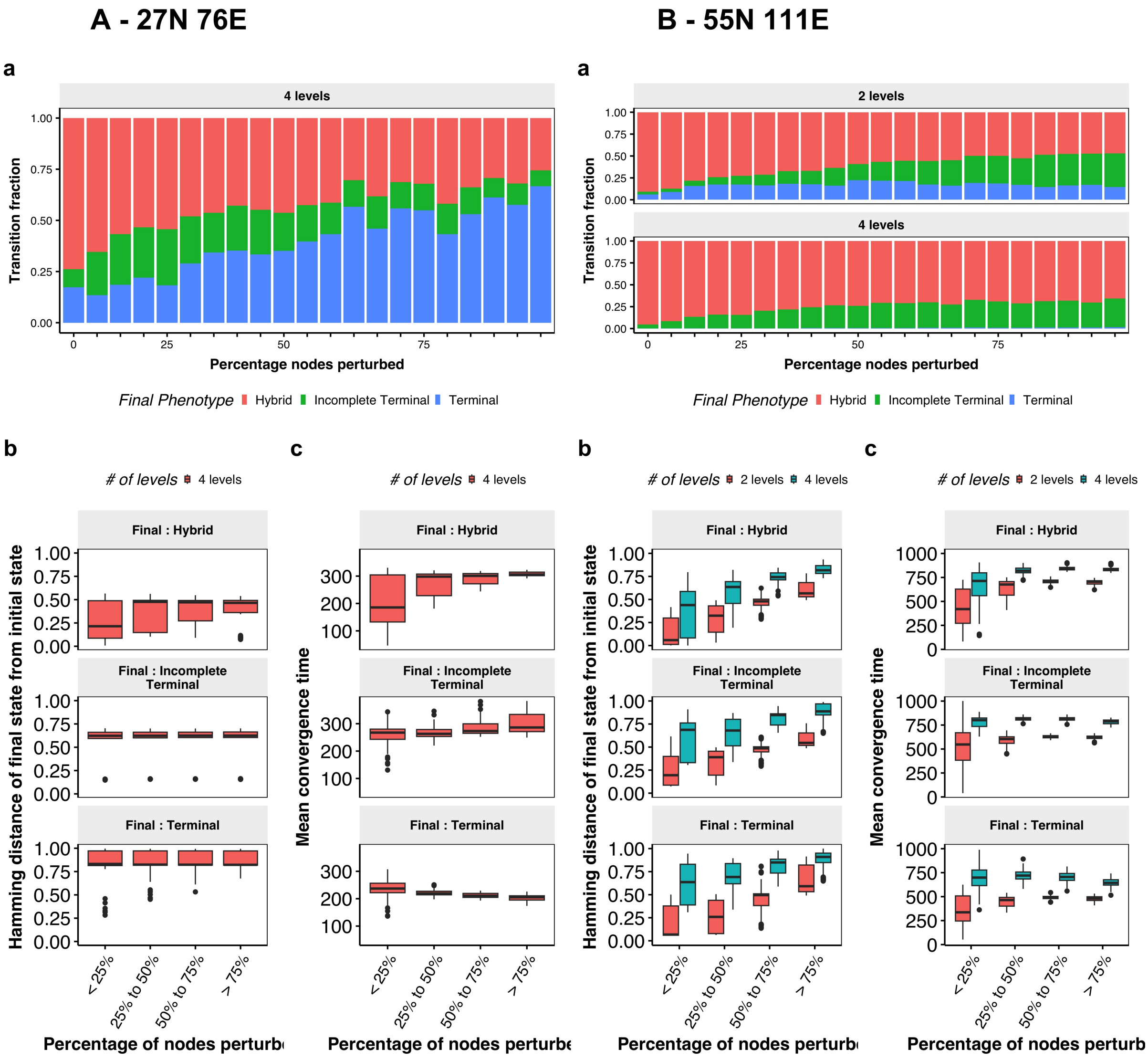


Figure S18: Analysis of multilevel model simulations for **A** 31N 95E[23] network and **B)** 57N 113E[21] network. The subpanel details are as follows: **a** Transition probability from hybrid states to other phenotypes for two and four-level formalism. Note that the increase in transition to hybrid phenotype in two-level model after 75% perturbation is an artefact of the symmetry of threshold-based formalism. **b** Boxplots depicting the hamming distance between the initial and final steady state, with each panel showing classified by the phenotype of the final state. **c** Same as b, but shows the time to converge.
